# Variation in reported human head tissue electrical conductivity values

**DOI:** 10.1101/511006

**Authors:** Hannah McCann, Giampaolo Pisano, Leandro Beltrachini

**Affiliations:** School of Physics and Astronomy, Cardiff University, Cardiff, UK; Cardiff University Brain Research Imaging Centre (CUBRIC), Cardiff, UK

**Keywords:** head conductivity, electrical impedance tomography, magnetic resonance electrical impedance tomography, electroencephalography, magnetoencephalography, electromagnetic source localisation

## Abstract

Electromagnetic source characterisation requires accurate volume conductor models representing head geometry and the electrical conductivity field. Head tissue conductivity is often assumed from previous literature, however, despite extensive research, measurements are inconsistent. A meta-analysis of reported human head electrical conductivity values was therefore conducted to determine significant variation and subsequent influential factors. Of 3,121 identified publications spanning three databases, 56 papers were included in data extraction. Conductivity values were categorised according to tissue type, and recorded alongside methodology, measurement condition, current frequency, tissue temperature, participant pathology and age. We found variation in electrical conductivity of the whole-skull, the spongiform layer of the skull, isotropic, perpendicularly- and parallelly-oriented white matter (WM) and the brain-to-skull-conductivity ratio (BSCR) could be significantly attributed to a combination of differences in methodology and demographics. This large variation should be acknowledged, and care should be taken when creating volume conductor models, ideally constructing them on an individual basis, rather than assuming them from the literature. When personalised models are unavailable, it is suggested weighted average means from the current meta-analysis are used. Assigning conductivity as: 0.41 S/m for the scalp, 0.02 S/m for the whole skull, or when better modelled as a 3-layer skull 0.048 S/m for the spongiform layer, 0.007 S/m for the inner compact and 0.005 S/m for the outer compact, as well as 1.71 S/m for the CSF, 0.47 S/m for the grey matter, 0.22 S/m for WM and 50.4 for the BSCR.

## 1. INTRODUCTION

Understanding electrical activity propagation throughout the head is essential in neurophysiology. In particular, forward and inverse solutions for source reconstruction in electroencephalography [EEG (Beltrachini, 2019a,b)], magnetoencephalography [MEG (Haueisen *et al.*, 1997; Vorwerk *et al.*, 2014)], transcranial magnetic stimulation [TMS; (Opitz *et al.*, 2011; Salinas, Lancaster, & Fox, 2009)] and deep brain stimulation [DBS; (Butson *et al.*, 2007; Dabek *et al.*, 2016; McIntyre *et al.*, 2004)] are governed by such phenomenon. Accurate values of head tissue electrical conductivity are vital to model and localise primary current generators within the brain based on both invasive and non-invasive recordings. Misspecification of tissue conductivities can consequently contribute to significant errors in magnetic field strength and electric surface potential estimations (Cohen & Cuffin, 1983a; Haueisen *et al.*, 1995; Okada, Lahteenmaki, & Xu, 1999), which may additionally introduce systemic errors in the EEG and MEG forward problems (Goncalves, *et al.*, 2003; Goncalves *et al.*, 2003) and result in inaccurate source localisation (Akhtari *et al.*, 2002; Haueisen *et al.*, 2002; Pohlmeier *et al.*, 1997; Vatta, Bruno, & Inchingolo, 2002). Anwander and colleagues (2002), for example, revealed mean EEG source localisation errors of 5.1mm and 8.88mm for radially- and tangentially-oriented sources, respectively, if white matter (WM) anisotropy was neglected in conductivity models. Whilst Hallez and colleagues (2005) reported average errors of 11.21mm, increasing to 13.73mm if skull anisotropy in addition to WM anisotropy was not considered. Even accounting for skull anisotropy has yielded maximum localisation errors of 6mm (Dannhauer *et al.*, 2011). Miscalculations can further lead to incorrect and inappropriate conclusions made regarding brain function, pathology and disease treatments inferred from E/MEG data (Wendel, Malmivuo, & Ieee, 2006). Most notably regarding implications in epilepsy treatment (Akhtari *et al.*, 2006; Fabrizi *et al.*, 2006), brain stimulation (De Lucia *et al.*, 2007; Sadleir *et al.*, 2010; Suh, Lee, & Kim, 2012) and insights into psychiatric and neurological disorders (Frantseva *et al.*, 2014; Park *et al.*, 2002; Schlosser *et al.*, 2007).

Currently, head tissue conductivity values are often assumed from the literature to create a volume conductor model. Despite extensive research and subsequent review papers (Faes *et al.*, 1999; Gabriel, Gabriel, & Corthout, 1996a; Geddes, & Baker, 1967), considerable differences in conductivity are evident between and within reports. Head tissue segmentation is known to be of substantial importance when assigning conductivity values (Akhtari *et al.*, 2000), however there remains discrepancies between such segmentation, for example, consideration of the various layers of the skull (Akhtari *et al.*, 2002), the importance of the dura layer (Ramon *et al.*, 2014) and the influence of blood vessels on high resolution EEG head modelling (Fiederer *et al.*, 2016). Additionally, accounts are inconsistent for the influence of anisotropy on conductivity values (Güllmar, Haueisen, & Reichenbach, 2010; Nicholson, 1965) and the brain-to-skull conductivity ratio [BSCR; (Gutiérrez, Nehorai, & Muravchik, 2004; Wolters et al., 2006)]. Furthermore, existing reports of conductivity vary depending on participant demographics, such as age and pathology, as well as measurement condition (i.e. *in vivo, ex vivo* or *in vitro*), applied frequency, tissue temperature and employed methodology.

**Figure 1.**
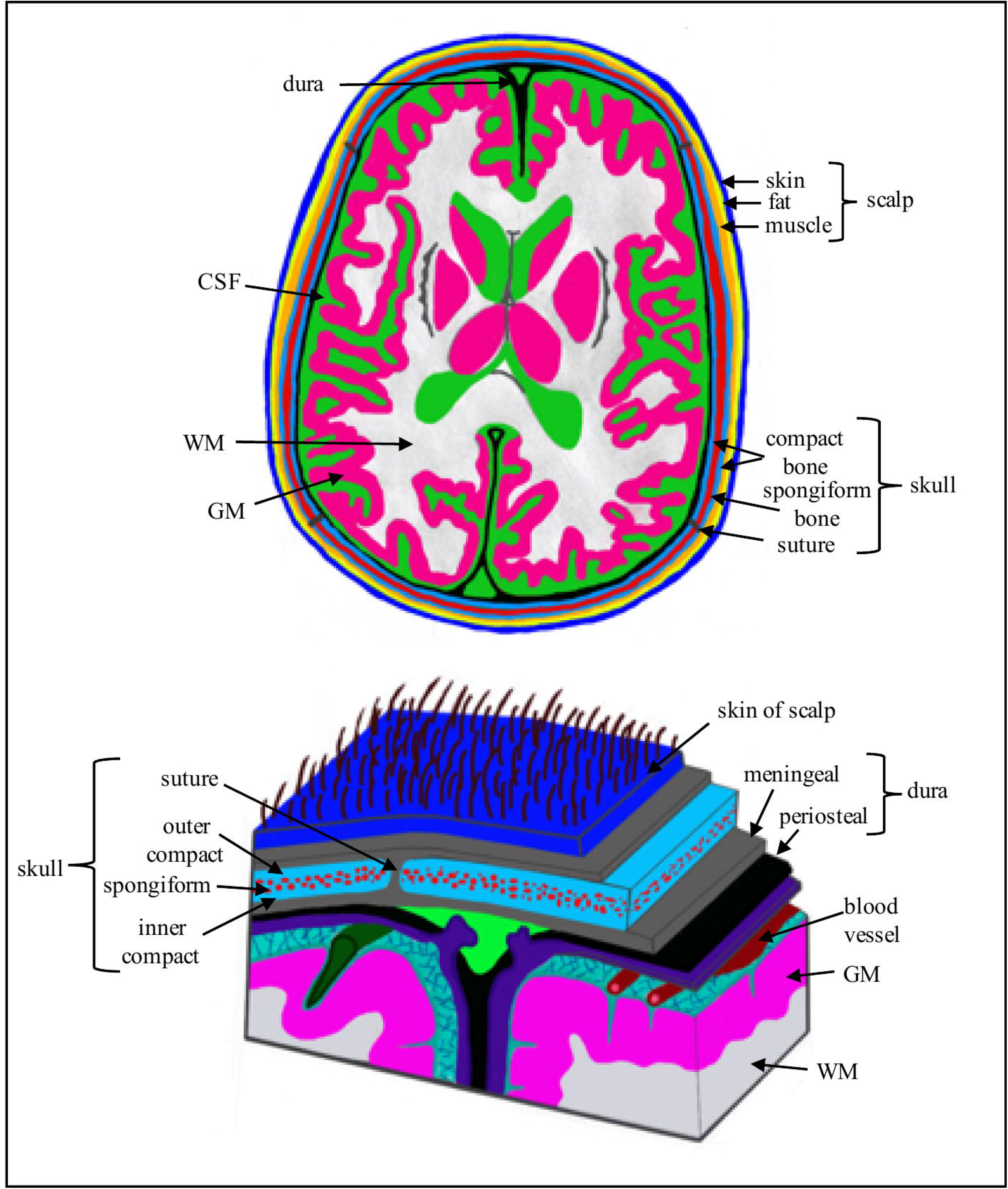
Figure displaying the various tissue compartments of the head and a subfigure of the detailed layers of the scalp, skull and brain.

Unsurprisingly, utilising different methodologies, such as directly applied current (DAC), electrical impedance tomography (EIT), E/MEG, Magnetic Resonance EIT (MREIT) and Diffusion Tensor Imaging (DTI), yield diverse conductivity values. The relative strengths and limitations of these methods is essential to accurately characterise discrepancies and inform future research. With DAC we refer to any invasive method where current is directly applied to tissue, either via multiple implanted electrodes or onto excised tissue, and electrical conductivity is determined from the resulting potential difference between a pair of electrodes. DAC methods have the advantage of not requiring a computational head model, which often introduces simplified assumptions regarding the neurobiology and dynamics of the human head, as well as being cost effective with a low acquisition time, easily portable and useable and has the potential to analyse conductivity of all tissue types. DAC methods however, are invasive, requiring post-mortem samples or excised tissues that are not under biophysically natural conditions. Tissues obtained post-mortem, for example, are subject to biochemical processes initiated by death, such as changes in ion mobility and cell membrane polarisation, which consequently affect conductivity (Opitz *et al.*, 2017). Opitz and colleagues (2017) importantly demonstrated, despite controlling for confounding variables (i.e. temperature), that live and post-mortem intracranial electrical fields significantly differed. Similarly, excised tissues undergo various extracting, preservation and holding procedures (i.e. saline soaked, time since excision, etc.) which can change the electrolyte concentration (Akhtari *et al.*, 2002) and hence influence conductivity. On the other hand, EIT, E/MEG, MREIT and DTI methods are non-invasive and occur *in vivo*, having the advantage of remaining under natural conditions. Additionally, EIT and E/MEG are both portable and cost effective with low acquisition times, compared to MREIT and DTI methods which are non-portable, more expensive and with high acquisition times, but EIT and M/EEG have lower spatial resolution than MREIT and DTI and require the use of a computational head model. Both MREIT and DTI however, employ magnetic resonance (MR) imaging, making skull conductivity non-accessible due to weak MR signal towards bone layers. A further advantage of DTI is the ability to classify anisotropic and heterogenous conductivity values of soft tissues (Johansen-Berg & Behrens, 2013). A summary of the strengths and weaknesses of the described methods are provided in Table 1.

**TABLE 1.**
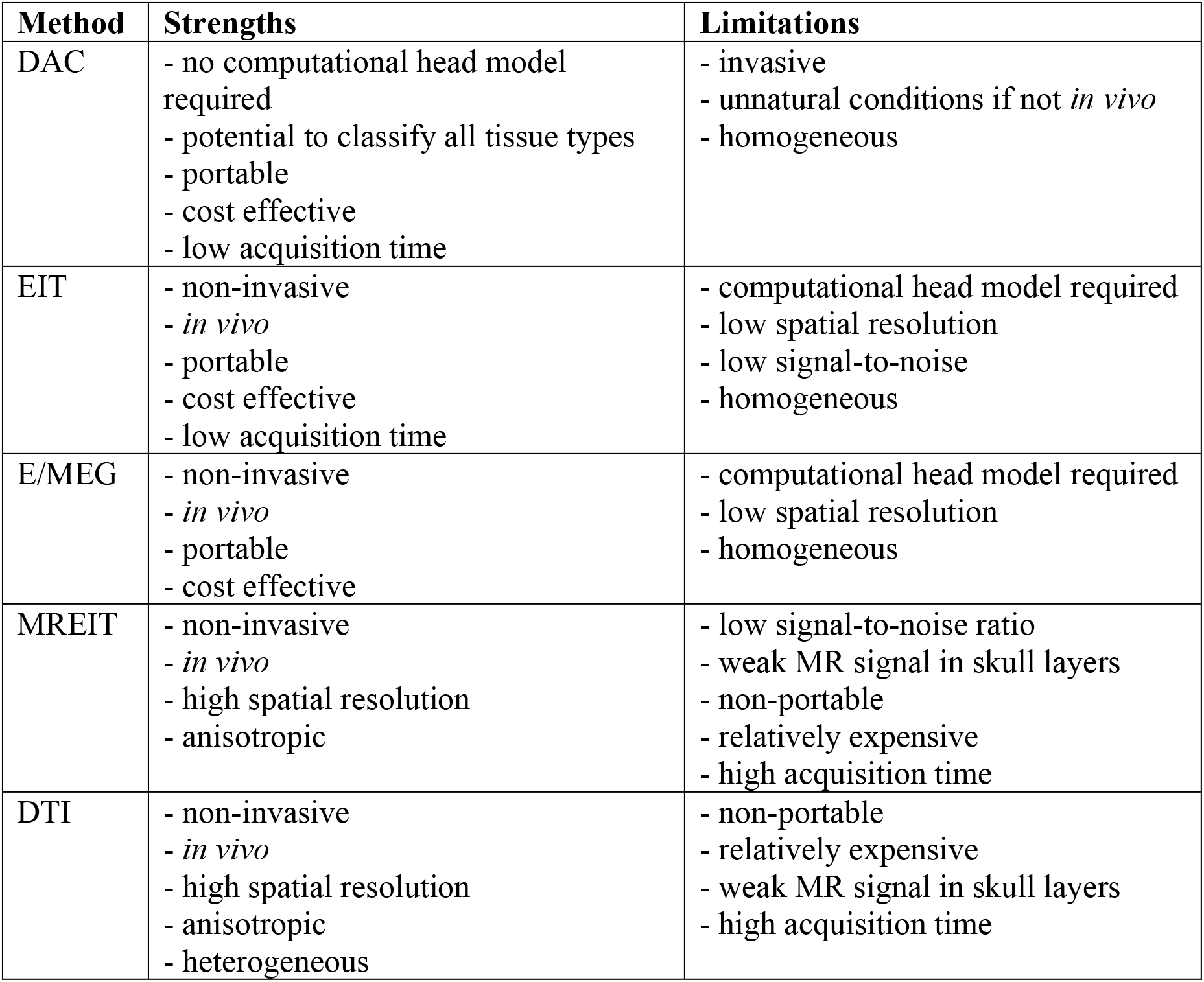
Methodology Strengths and Limitations

Considering the vast disparity in data, assuming conductivity from previous literature is insufficient when accurate and precise values are required. Significant and important factors affecting this variation, however, are currently unknown. Knowledge of influential variables, such as tissue segmentation, methodology employed, sample temperature or participant pathology can provide insights into the stability of tissue conductivity values and methodology, as well as suggest areas for future research. The current study aimed to systematically and extensively investigate all published reports of human head tissue electrical conductivity to i) evidence any significant variations in conductivity values of different head tissue types; ii) determine any significant factors contributing to variation; and iii) analyse the impact these factors may have on source reconstruction in E/MEG. A systematic review, restricted to human head tissue, was carried out to identify relevant papers, and a meta-analysis was completed to reveal significant factor variables via a multiple regression. It was hypothesised that head tissue conductivity would vary between and within tissues. It was expected the meta-analysis would further reveal significant influential factors and their impact.

## 2. METHODS

### 2.1 Literature search

Preferred Reporting Items for Systematic reviews and Meta-Analyses (PRISMA) statement guidelines (Moher *et al.*, 2009) were followed and a PRISMA checklist and flow diagram were completed (Appendix A.) An extensive literature search, spanning three databases (PubMed, Scopus and Web of Knowledge), was conducted to retrieve published and peer-reviewed studies exploring electrical conductivity (or equivalent) of the human head (or equivalent). The keywords utilised for the systematic literature search are provided in Appendix B. Article titles were systematically searched using relevant and/or equivalent keywords, unrestricted by year of publication, language or design. Reference lists of included papers were hand-searched to identify additional papers. Duplicates following the initial literature search were removed.

### 2.2 Selection criteria

Papers met the inclusion criteria if they i) provided at least one conductivity measure (or equivalent from which conductivity could be calculated), of the ii) human iii) head, where iv) employed methodology and v) tissue type were available. Reviews were only included as an information source to the original reference, where data was thus extracted. Exclusions were made if any of the five inclusion criteria were absent or ambiguous, or if an English version was unavailable after extensive search. In addition to conductivity value, methodology and tissue type, reports were collected on measurement condition (i.e. *in vivo, ex vivo*, or *in vitro*), applied frequency to determine the conductivity, tissue temperature, as well as participant’s age, gender and pathology. Missing information for one or more of these variables did not result in exclusion. Studies applying frequencies above 1KHz were excluded from analysis on the grounds this frequency is besides the scope of typical brain activity recorded in E/MEG.

All identified titles resulting from the literature search, following removal of duplicates, were initially screened for applicability and/or immediate exclusion. Remaining abstracts were further assessed, and full texts of potentially relevant papers were obtained to determine if they consequently met the inclusion criteria.

### 2.3 Data extraction and synthesis

All conductivity, resistivity or impedance values were extracted from each paper and converted to S/m for standardisation. The sample mean and standard deviation were subsequently calculated for every differentiation in methodology within each paper and characterised according to the aforementioned variables.

### 2.4 Variable Definitions and Classification

#### a) Tissue Types

For the current review, tissues were separated into four major compartments, each comprised of sub-compartments: the scalp (*skin, fat, muscle*), the skull (*spongiform, inner* and *outer compact bone* and *sutures*), cerebrospinal fluid (*CSF*) and the brain [grey matter (*GM*), *WM*, the *dura* layer, *blood, cerebellum, lesions*, epileptogenic zone (*EZ*)]. Conductivity values were assigned according to tissue type as reported. Tissues were classified as *whole-scalp*, *whole-skull* or *whole-brain* when no conductivity values for their sub-compartments were reported, similarly *whole-compact* bone was assigned if no values for the inner and outer compact bone were provided. If given, WM was further segmented into WM oriented in parallel (*WM_par*) or perpendicular (*WM_perp*) to the applied current. See Figure 1 for a detailed representation of all tissue compartments. Additionally, when available, the *BSCR* was reported as a nominal ratio without units.

#### b) Measurement conditions

Conditions were separated into three main categories:

*In vivo*–“within the living”; experiment conducted on or in whole living organisms/cells. Electrical conductivity values obtained within a living head were considered *in vivo*.

*Ex vivo*–“out of the living”; experiment in or on tissue from an organism in an external environment, but with minimal alteration of natural conditions, e.g. cultured cells derived from biopsies. Experiments where tissue was excised but kept within conditions similar to the human head were characterised as *ex vivo*.

*In vitro*–“within the glass”; experiment within a controlled artificial environment outside of a living organism, isolated from their usual biological surroundings e.g. in a test tube/dish. Measurements where tissue was excised and stored in environments unlike the human head were classified as *in vitro*.

#### c) Measurement methods

Data acquisition techniques were categorised into five groups:

*DAC* - invasive method of determining electrical conductivity, where a current was directly applied to the tissue, either via implanted electrodes in the head, or onto excised samples. The resulting electric potential difference from the applied current is measured via additional (implanted or applied to excised tissue) electrodes to calculate the electrical conductivity. Studies where electrical current was directly and invasively applied to the head tissue were characterised as DAC.

*EIT* – a non-invasive medical imaging technique where alternating current at single or multiple frequencies is applied to the skin through two or more conducting surface electrodes. The resulting potential difference between the remaining measuring electrodes is then recorded. From this the electrical conductivity, permittivity and impedance can be inferred to create a tomographic image (Barber & Brown, 1984; Henderson & Webster, 1978). Papers indicating an applied current of less than 1kHz, injected through any number of electrodes and the resulting voltage were classified as EIT.

*MREIT* – measures the induced magnetic flux density resulting from an injected current (as in EIT) using a magnetic resonance imaging (MRI) scanner. The internal current density is then computed and combined with magnetic flux density measurements to perform conductivity map reconstruction, using various inverse solutions (Bodenstein, David, & Markstaller, 2009). Studies specifying acquisition of MRI data during current injection (as in any EIT method) to reconstruct conductivity (using any inverse method) were categorised as MREIT.

*E/MEG* – electromagnetic data recorded from E/MEG employed to iteratively estimate the equivalent electrical conductivity that best matches the computed source localisation given the obtained E/MEG data (Baysal and Haueisen, 2004). Articles estimating conductivity by employing data from E/MEG (of any set up) were characterised as E/MEG.

*DTI* – diffusion-weighted MR images of the brain are acquired to measure the diffusion tensor eigenvalues, from which the electrical conductivity tensor eigenvalues are directly calculated (Sekino, Inoue, & Ueno, 2005; Tuch *et al*., 1999; Tuch *et al.*, 2001). Texts using diffusion imaging (of any protocol) to explicitly estimate the electrical conductivity tensor map were considered as employing DTI methodology for the current review. This included any method for estimating conductivity from the diffusion tensor. DTI papers where conductivity was not explicitly reported were not included in the current review.

#### d) Frequency

Frequency of applied or injected current (if applicable). Frequency was not extracted from papers where this was not specified.

#### e) Temperature

Classified according to whether the tissue sample was measured at/near *body* temperature (37°C) or *room* temperature (18-25°C). Unknown values were not reported for analysis.

#### f) Participant’s Age

When available, mean and standard deviation of participant’s age were calculated for each paper and recorded in Table 2. Age at time of death was recorded for deceased participants. If specific age was unavailable, age was characterised as *adult* (all participants were over the age of 18), *paediatric* (all participants were under the age of 18), or *both* (participants were a mixture of over and under the age of 18).

#### g) Participant pathology

Participants were characterised as *healthy* if they had no neurological, developmental or psychological deficits, as reported in the research paper. Pathology was categorised as *epilepsy* for studies recruiting patients that presented with any classification of epileptic seizure. Similarly, *tumour* was assigned to papers where patients displayed any type of tumour, and *neuro* to patients with any type of neurological disorder that was not otherwise classifiable. Further pathologies included *Parkinson’s Disease, Alzheimer’s Disease* and *stroke*. All conductivity values were assumed to originate from healthy tissue, within the classified pathology, unless otherwise stated. Pathology was reported as unknown if not available in the literature.

### 2.5 Quality Analysis

Drawing robust conclusions from systematic reviews and meta-analyses requires consideration of the systematic and random errors introduced in each included study by “assessing the methodological quality” (Moher, Jadad, & Tugwell, 1996; Verhagen *et al.*, 2001) in order to estimate “risk of bias”. Various tools are available for assessing study quality and addressing the systematic errors in each study, however none specifically to assess the quality of studies measuring the electrical conductivity of the human head. The current meta-analysis therefore, made use of the Cochrane Collaboration recommended Quality Assessment of Diagnostic Accuracy Studies (QUADAS) checklist (Whiting *et al.*, 2003), where each item was adjusted for relevance, and any additional relevant items were added. A scaled numerical value was further assigned according to the studies compliance with each item; any irrelevant items were ignored. The sum, divided by the number of relevant items, was subsequently calculated to provide a final Quality Assessment Score (QAS), with an absolute maximum value of one (the closer the score is to one, the more reliable the study was considered). To ensure reliability of the QAS’s, papers were chosen at random and QASs calculated by two researchers, any discrepancies were discussed and if not resolved the mean QAS was assigned. The employed Quality Assessment Protocol and three examples are provided in Appendix C.

In addition to accounting for systematic errors within each study, random errors produced from inherently unpredictable variation in methodology were accounted for. This was adapted from the guidelines provided by Rosenthal (1991) and Borenstein and colleagues (2011) for meta-analysis weighting. Confidence values for each measurement were calculated to indicate the confidence each value of conductivity was 100% accurate. Firstly, the relative error was calculated for each conductivity value, as the standard deviation percentage of a multitude of values for a single tissue type for each participant (if the method is 100% precise, each value for the same tissue should be the same) or the error attributed to the measurement protocol – both described as a decimal. If both the standard deviation and measurement error were provided, the standard deviation was used to calculate the relative error. The relative error was then subtracted from one (where one indicates complete confidence the conductivity value is 100% accurate) to obtain a final confidence value of which the maximum is one. For example, a reported conductivity value with an associated standard deviation percentage of 8% will receive a confidence value of 0.92. Alternatively, when the standard deviation was not provided, the experimental error was utilised instead; e.g. a study with a methodological error of 0.05 would receive a confidence value of 0.95.

To incorporate both the systematic and random errors associated with each study, the Quality Assessment Score of each study and the confidence values of each conductivity value were combined to provide a “weight”. This weight was calculated by multiplying the QAS by the confidence value (both with a maximum of one). The maximum associated weight each value has towards the analysis is therefore one. Values assigned weights closer to one were therefore regarded as being more accurate.

### 2.6 Statistical analysis

Data was pooled and grouped according to tissue type, in order to determine i) the variation in conductivity for each tissue, ii) which significant variables account for differences in conductivity, iii) whether mean conductivity values for each tissue type are statistically different depending on employed methodology and participant demographics, and iv) reveal any statistical relationship between conductivity and reported variables.

Boxplot diagrams, presenting the range, median and mean of conductivity measurements for each tissue type were created to demonstrate variation in conductivity within different tissues. For each tissue with more than three results in at least two variables, a weighted multiple regression was carried out using SPSS (Corp, 2013). The dependent variable (conductivity) was regressed against every independent variable (IV; measurement condition, method, frequency, temperature, age and pathology) collectively, to determine the proportion of variance accounted for by all factors, and individually to discover significant factors predicting variation in conductivity. Weights for each conductivity value were assigned according to the Quality Analysis described above (section 2.5). A two-tailed t-test (when comparing two independent variables) or a one-way Analysis of Variance (ANOVA; when comparing more than two independent variables) was conducted to reveal differences in conductivity for each tissue, according to categorical IV’s previously revealed to account for a significant proportion of variance. A Pearson correlation analysis was alternatively conducted for continuous IV’s accounting for significant variation to reveal any statistical relationships.

## 3 RESULTS

### 3.1 Search results

Following removal of duplicates, 3121 studies were identified through the literature and reference list search, of which 382 abstracts were screened for relevance and 211 full text articles were obtained and assessed for eligibility. A total of 155 papers were excluded (see Appendix A, Figure A.1: PRISMA flow diagram).

### 3.2 Included studies

A total of 56 studies (407 participants) were included in the quantitative synthesis (Table 2). Seventeen different tissue types were identified, using 5 methodologies and 3 measurement conditions. Conductivity was measured *in vivo* in 42, *in vitro* in 7 and *ex vivo* in 8 research papers. Measurements were obtained using DAC in 14 studies, using EIT in 10, E/MEG in 7, MREIT in 15 and DTI in 9 papers. Conductivity was acquired at frequencies varying between 0Hz and 1005Hz, and tissue temperatures between 18.5 and 37.5°C. Of the 23 articles that specified, total participant age ranged from 4 months to 87 years old, whilst the remainder classified subjects into adults or children. Forty papers reported on healthy participants, participants from 10 studies were diagnosed with epilepsy, patients with tumours were included in 3 studies, stroke patients were employed in 2 papers, whilst separate papers included patients with various neurological disorders and Parkinson’s Disease. Descriptive statistics for each tissue type are provided (Table 3), in addition to a boxplot displaying variation in conductivity values for different tissue types (Figure 2). The average mean was calculated for each tissue type, where all conductivity values contributed equally to the mean. A weighted average mean was additionally calculated to take into consideration the quality of each study and provide a recommended value that was obtained under suitable and realistic conditions. The weighted average mean and standard deviation (in S/m) for the main tissue types were: scalp = 0.41± 0.18, whole skull = 0.02 ± 0.02, spongiform skull layer = 0.048 ± 0.07, whole compact skull layer = 0.005 ± 0.002, outer compact = 0.005 ± 0.003, inner compact = 0.007 ± 0.004, CSF = 1.71 ± 0.3, GM = 0.47 ± 0.24, WM = 0.22 ± 0.17, BSCR = 50.4 ± 39. A boxplot evidencing the average weights assigned to each study according to the employed methodology is further demonstrated (Figure 3). Average study weights were revealed to be significantly different depending on methodology [F (4, 56) = 3.121, p=.022)].

**TABLE 2.**
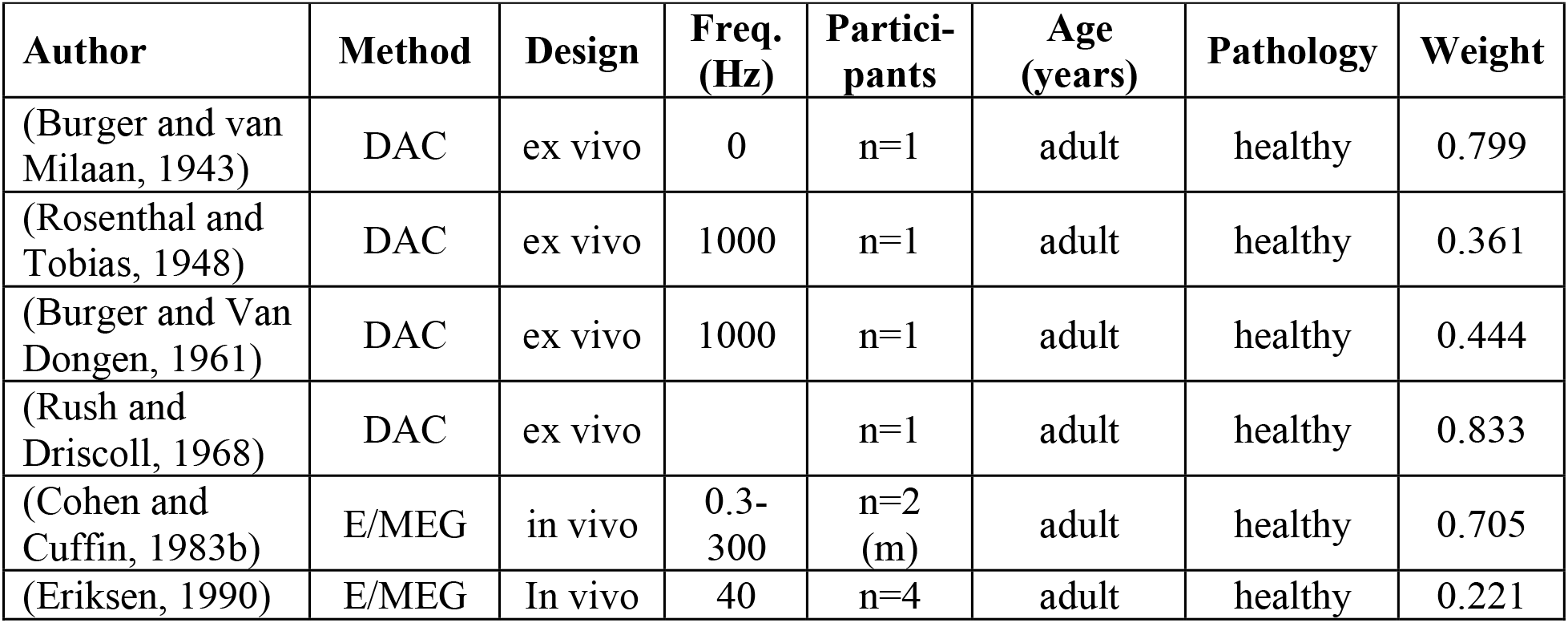
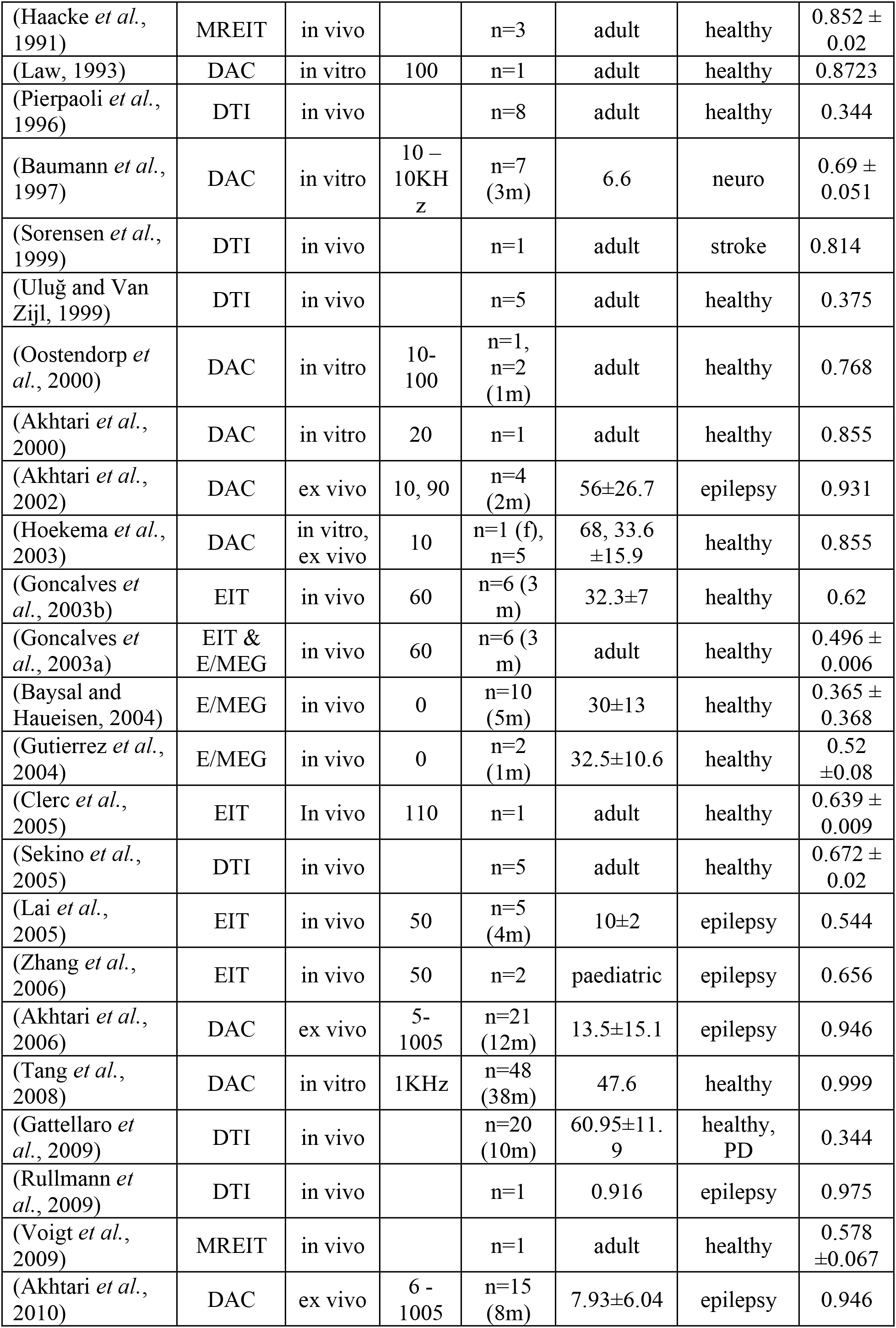
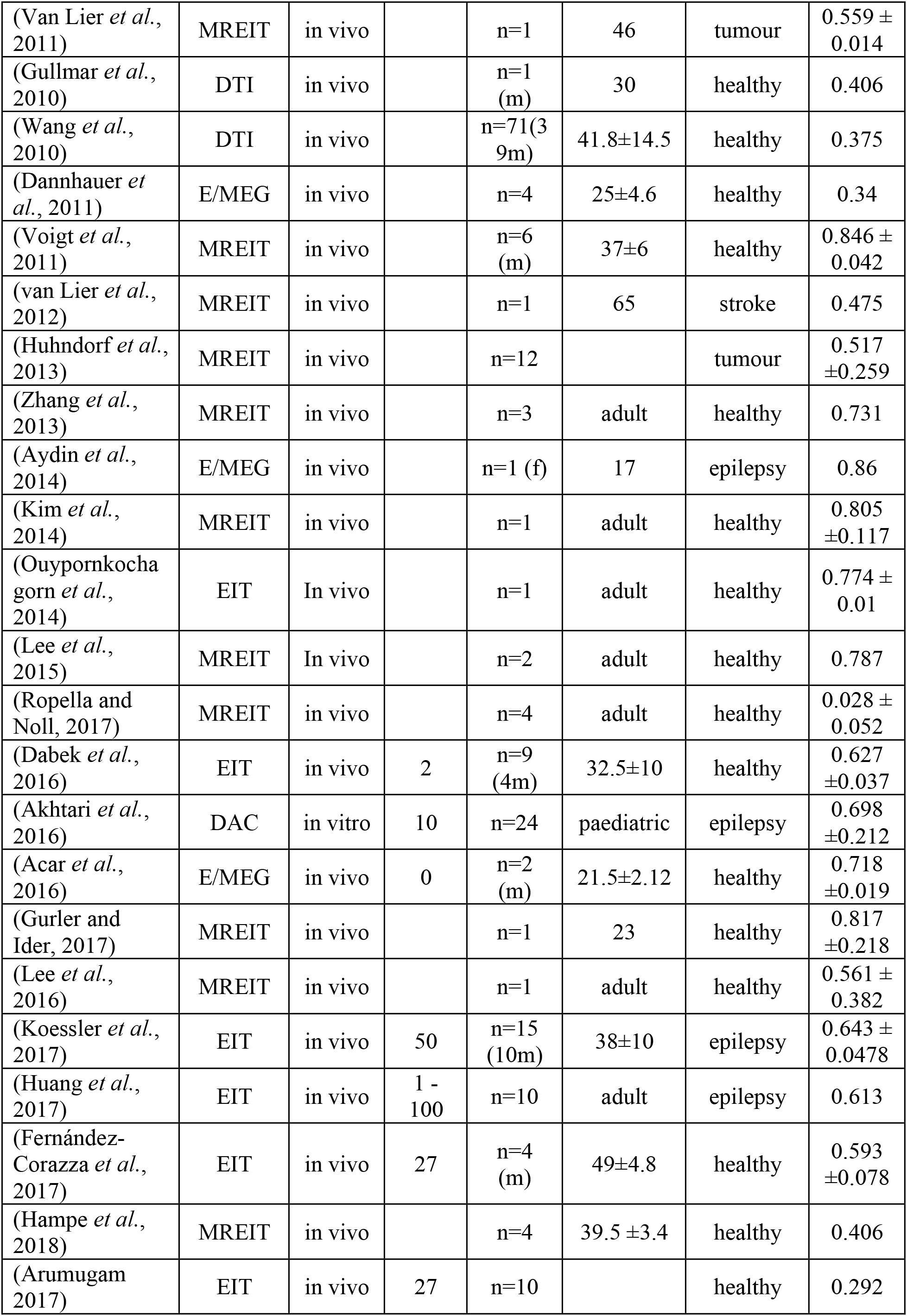
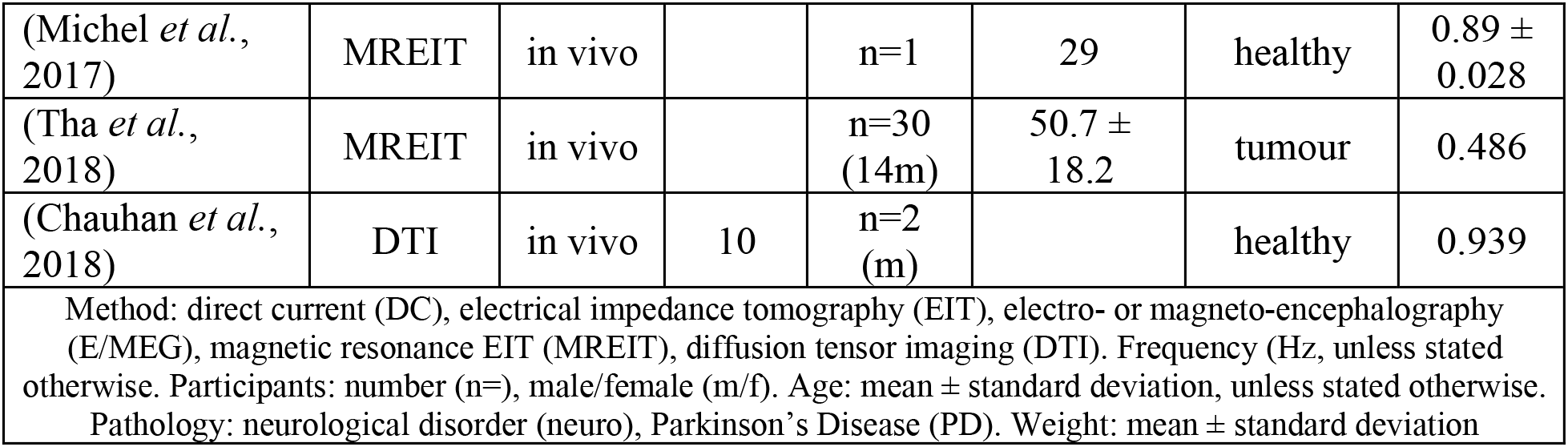
Summary of papers included in meta-analysis

**TABLE 3.**
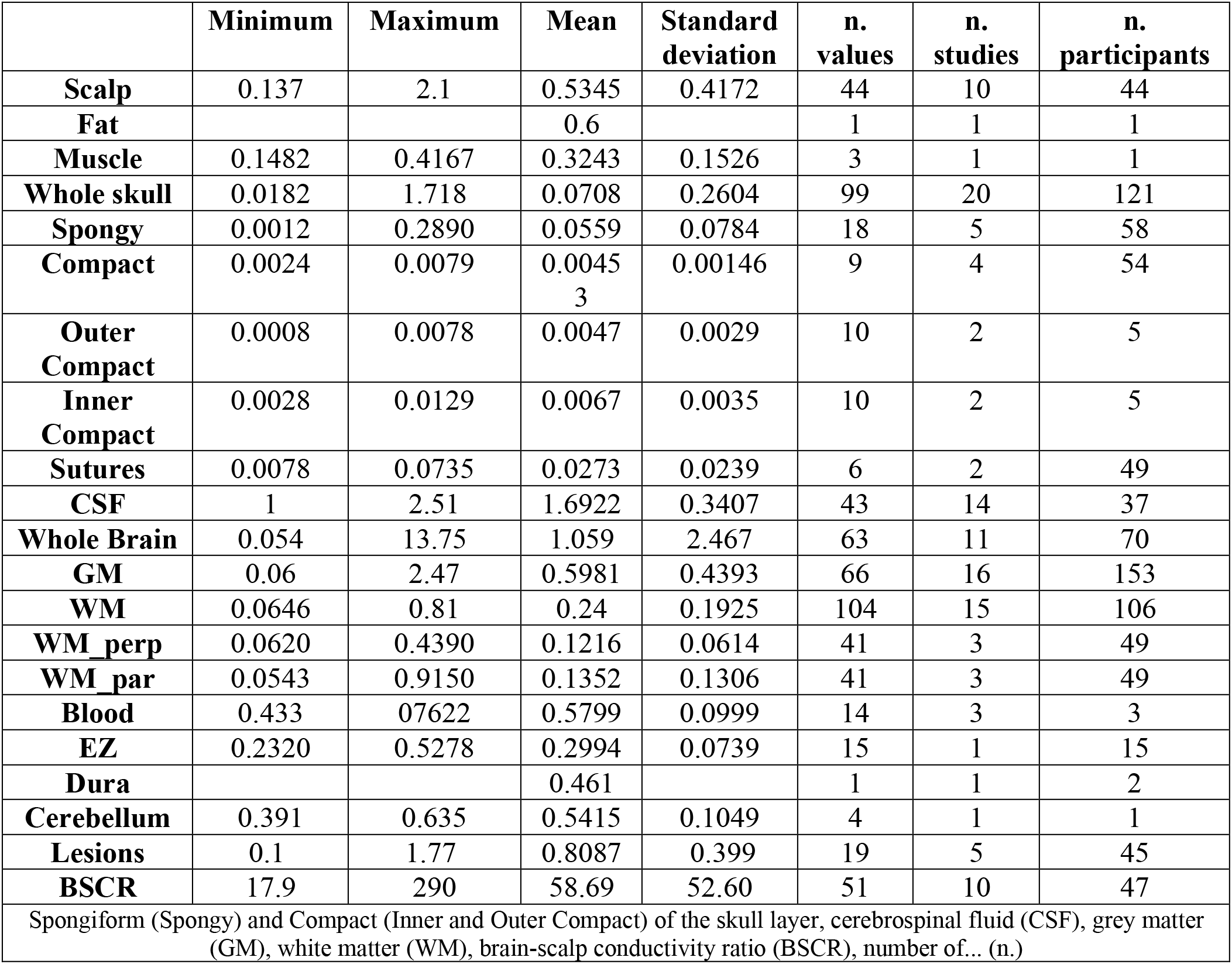
Descriptive statistics for each tissue type

**Figure 2.**
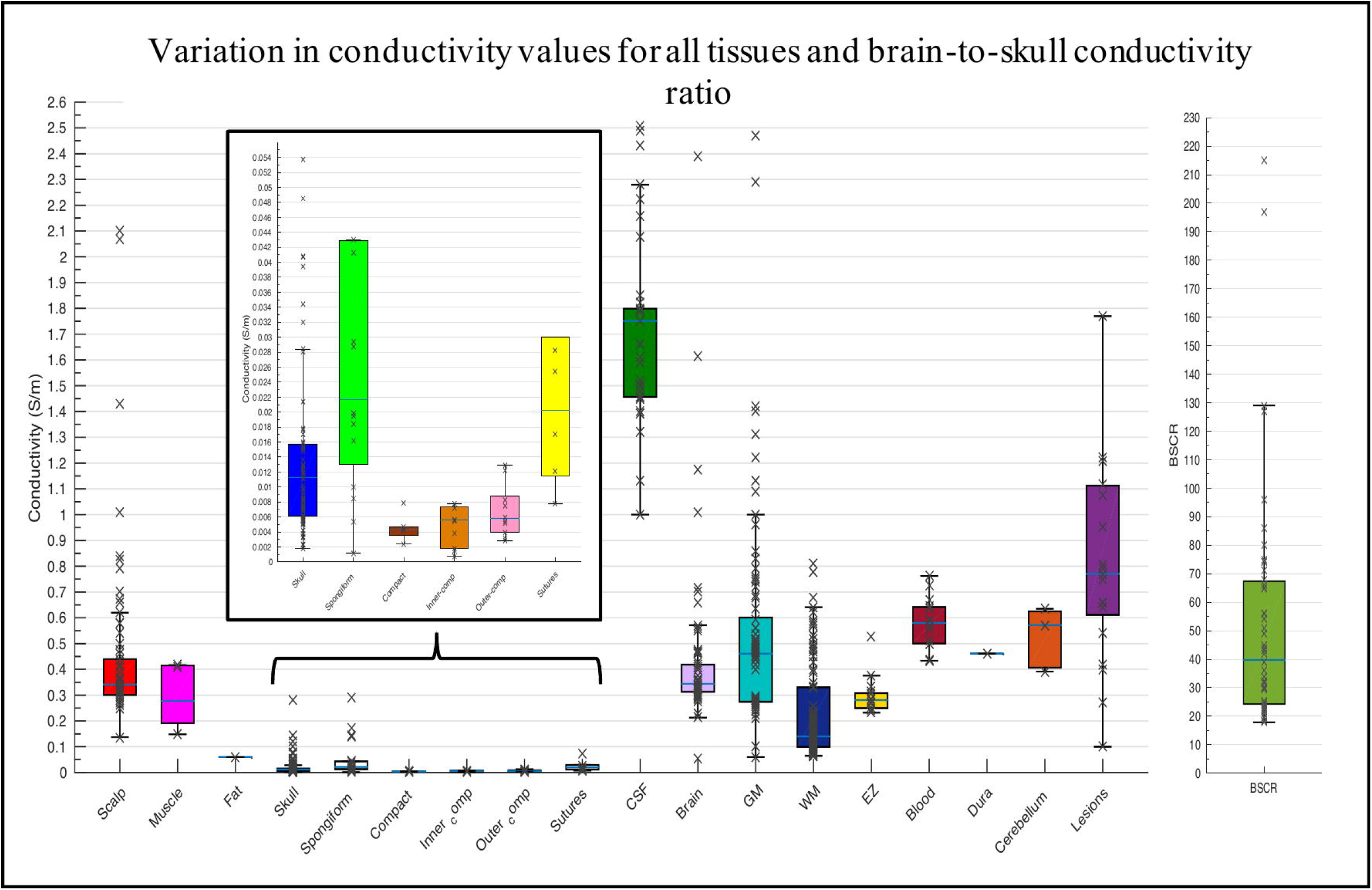
Boxplot displaying the inter-quartile range (first to third quartile as solid box), the median (solid blue line), minimum and maximum (solid whiskers) of all available conductivity values (S/m) for all tissues and BSCR displayed as crosses.

**Figure 3.**
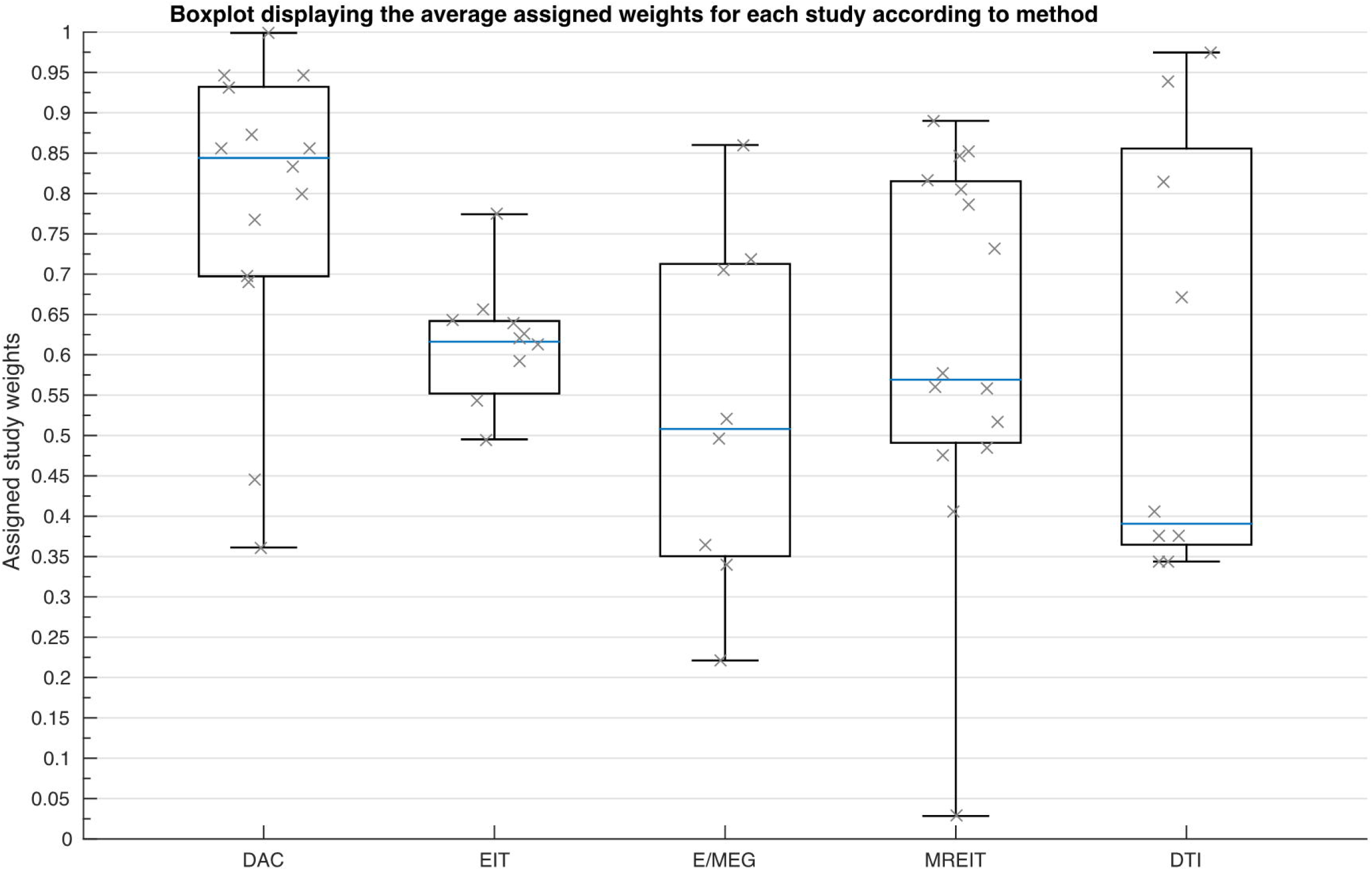
Boxplot displaying the mean value of the assigned weights for each study (indicated by a cross) dependent on the employed methodology.

Following visual inspection, it can be seen conductivity values vary considerably within and between tissue types. Insufficient data was available to calculate regression statistics for muscle, fat, blood, the epileptogenic zone, the dura layer and the cerebellum.

### 3.3 Scalp

A weighted multiple regression revealed scalp conductivity variation was insignificantly predicted by the IV’s collectively (p>.05). Although insignificantly different, a comparison between employed method (as shown in Figure 4) was made to graphically display any elevated values and further demonstrate variation despite statistical insignificance. Figure 4 further reveals less deviation within values for EIT than for E/MEG. Huang and colleagues (2017) yielded conductivity measurements significantly above the inter-quartile range. Additionally, Baysal and Haueisen (2004) revealed highly elevated conductivity values beyond the axis range displayed in Figure 4, with standard deviation >5000% (see Section 4.1 for further explanation of outliers).

**Figure 4.**
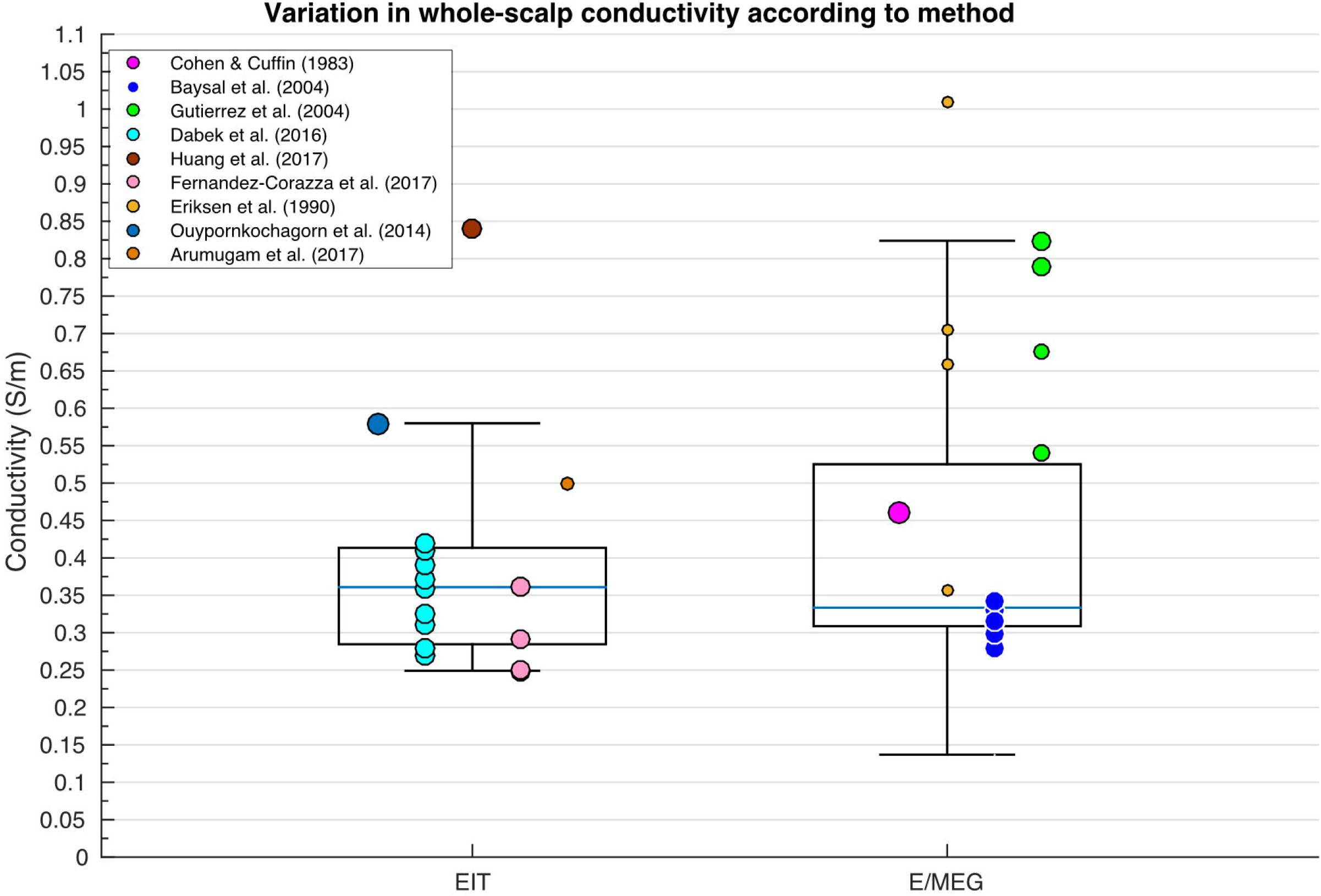
Boxplot displaying inter-quartile range (box), medium (solid blue horizontal line), maximum and minimum (upper and lower whiskers respectively) of scalp conductivity according to method for each available paper. Size of data points indicates relative weight of value.

### 3.4 Skull

#### 3.4.1 Whole-skull

A weighted multiple regression revealed deviation in whole-skull conductivity could be significantly predicted by the methodology, condition, temperature, frequency, pathology and age collectively [R^2^(6, 36) = .827, p <.001]. A one-way ANOVA revealed conductivity of the whole skull varied significantly according to employed methodology [Figure 5; F(2, 96) = 4.088, p=.020]. Differences in conductivity values for the whole-skull were statistically different according to method, where values obtained using EIT were significantly lower than those obtained with DAC and E/MEG.

**Figure 5.**
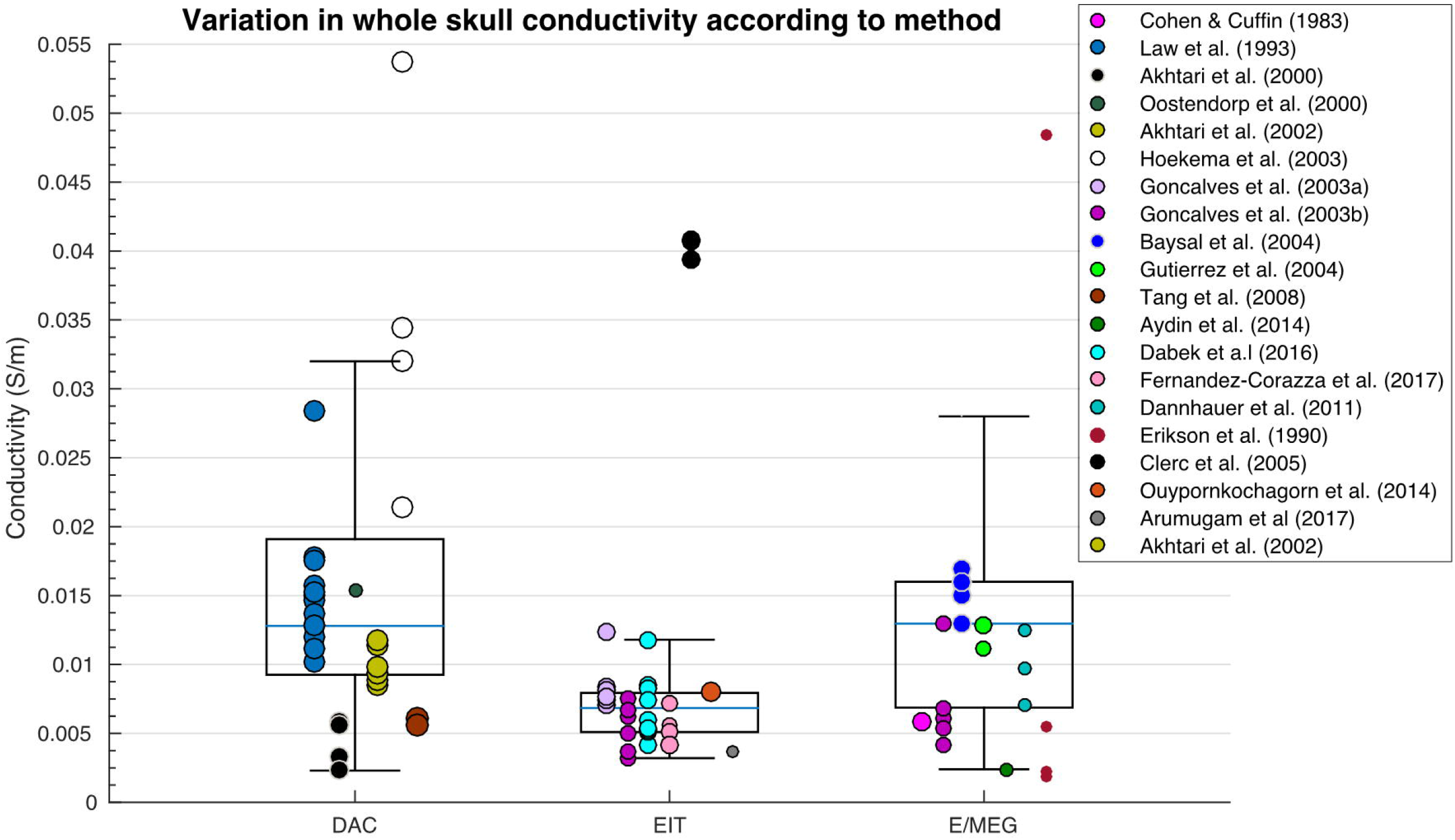
Boxplot displaying variation in whole-skull conductivity according to method.

#### 3.4.2 Spongiform bone skull layer

A weighted multiple regression revealed variation in conductivity values of the spongiform bone layer of the skull was significantly predicted by condition, temperature, frequency, pathology and age [R^2^(5, 6) = .832, p =.026]. Spongiform conductivity measurements were significantly different according to condition [Figure 6; F(2, 15) = 11.357, p=.001] and temperature [t(16)=2.449, p=.001).

**Figure 6.**
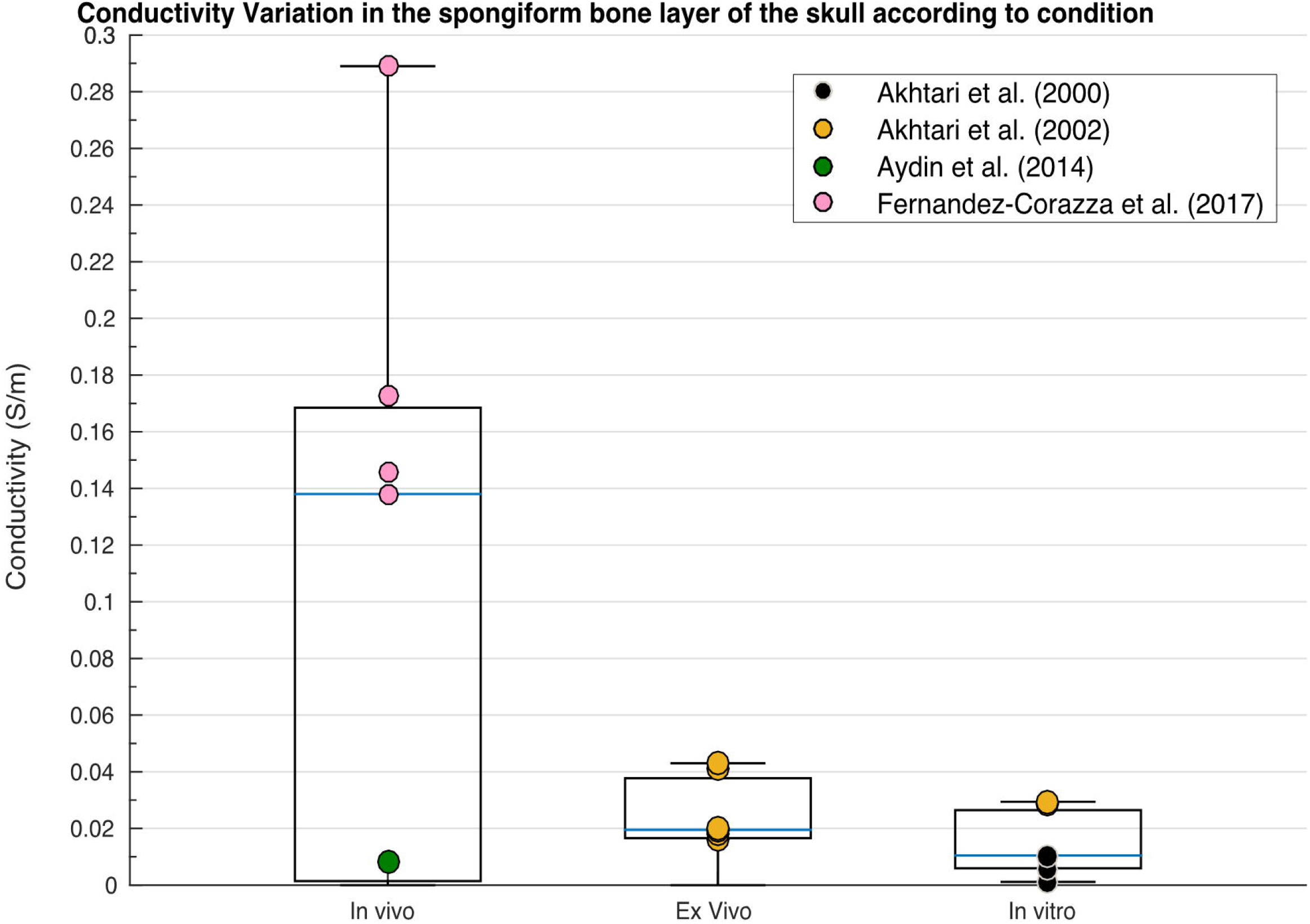
Boxplot displaying variation in conductivity of the spongiform bone layer of the skull according to condition.

#### 3.4.3 Compact bone skull layer

None of the IV’s significantly predicted variation in conductivity values of the whole compact layer, the inner compact bone layer or the outer compact bone layer according to the weighted multiple regression analysis. Despite insignificant results, a graphical representation of conductivity for the different compact bone layers revealed clear diversions within and between each of the layers (Figure 7).

**Figure 7.**
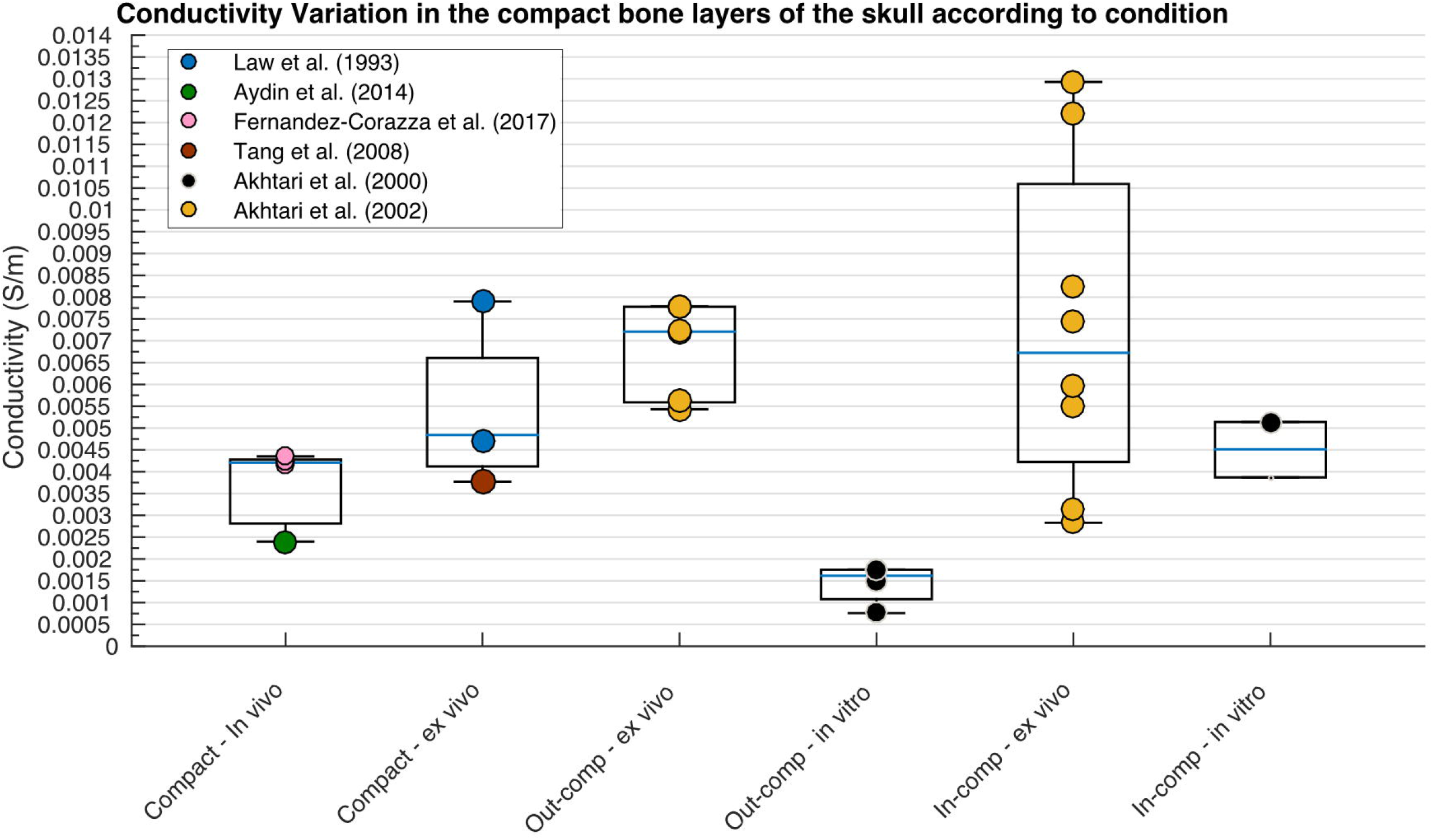
Boxplot displaying variation in conductivity of the compact layers according to condition.

### 3.5 Cerebrospinal Fluid (CSF)

Significant differences [t(36) = 2.695, p=.006] in measurements obtained at body (~1.79S/m) and room (~1.45 S/m) temperature as revealed from Baumann and colleagues (Baumann *et al.*, 1997), hence values at room temperature were revealed prior to regression and comparison of means analysis. Variability in CSF conductivity was discovered to be insignificantly explained by the weighted multiple regression model. Despite insignificant results, the boxplot in figure 8 allowed for a graphical representation of the large spread of values obtained using MREIT.

**Figure 8.**
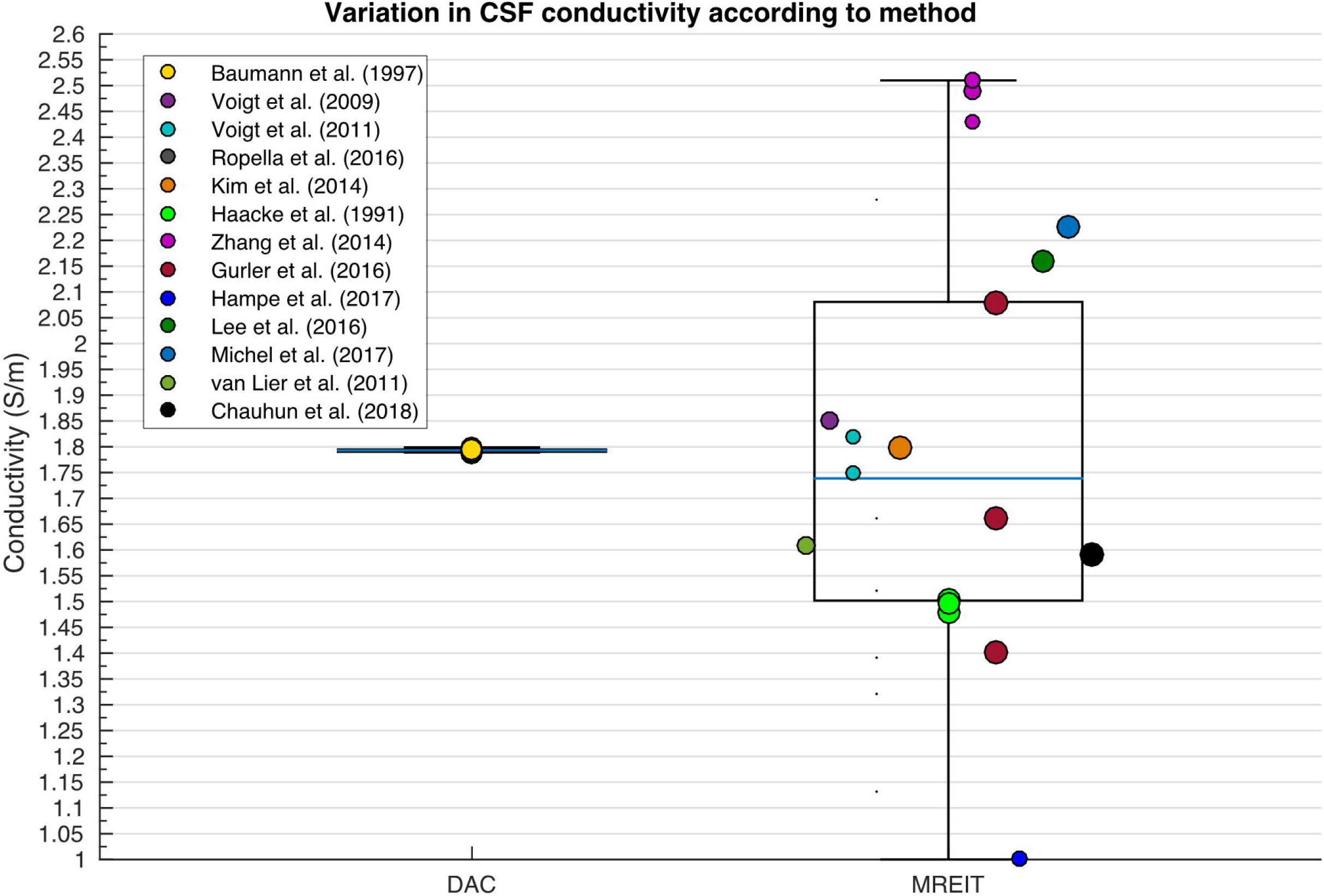
Boxplot displaying variation in CSF conductivity depending on method.

### 3.6 Brain

Differences in whole-brain conductivity values were not significantly predicted by the independent variables according to the weighted multiple regression analysis. Figure 9 reveals the large variation in data obtained for conductivity values of the whole-brain for each methodology, suggesting no one method generates a stable result for conductivity of the brain as a whole compartment.

**Figure 9.**
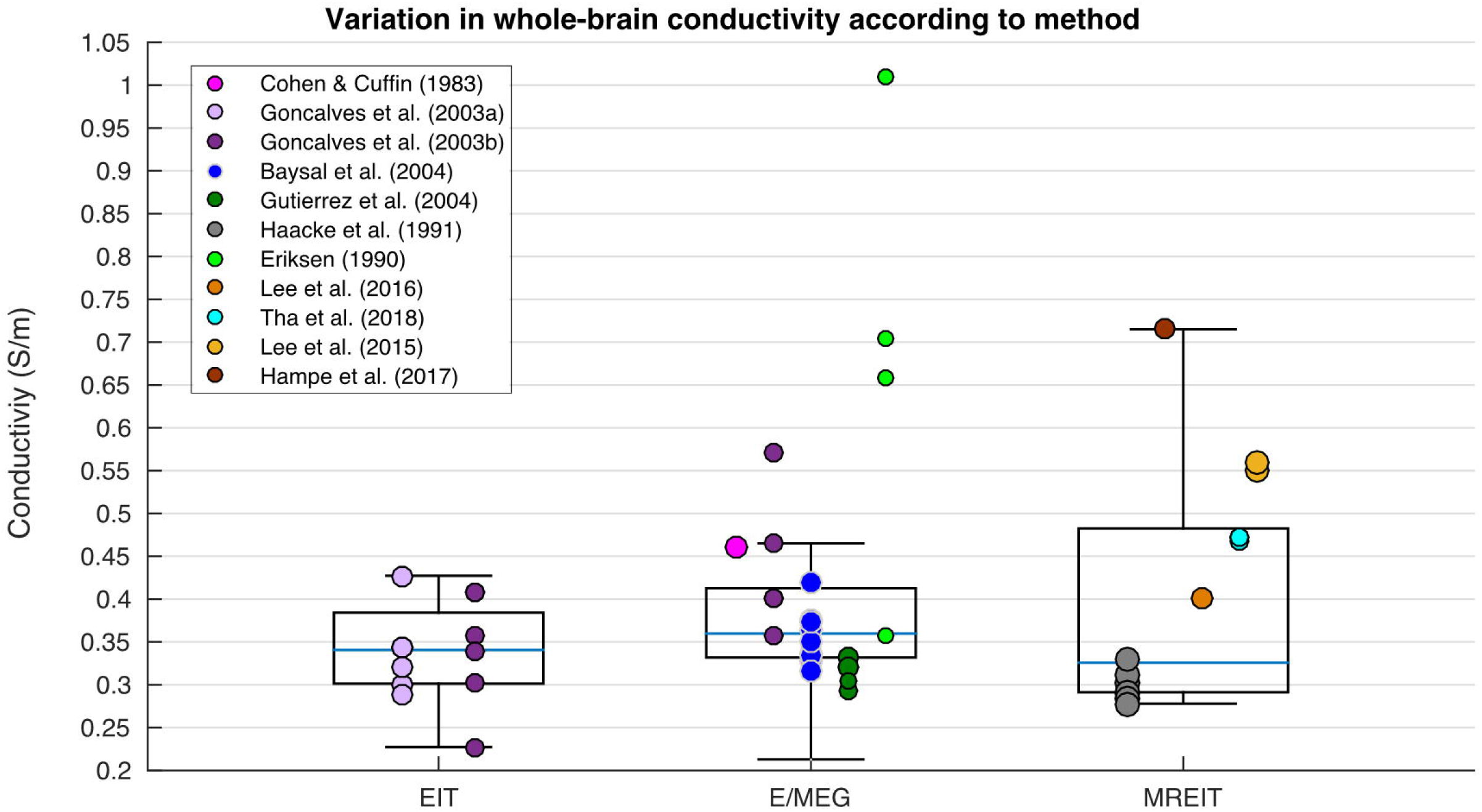
Boxplot displaying variation in whole-brain conductivity depending on method.

#### 3.6.1 Grey Matter

Variation in GM conductivity was not significantly explained by the weighted multiple regression model. However, a one-way ANOVA determined significant differences in GM conductivity according to method [Figure 10; F(3, 62) = 17.896, p<.001], where results obtained with MREIT were significantly higher than DTI which were in turn significantly higher than EIT. Pathology further yielded significantly different conductivity results for GM [Figure 11; F(4, 61) = 2.968, p=.026].

**Figure 10.**
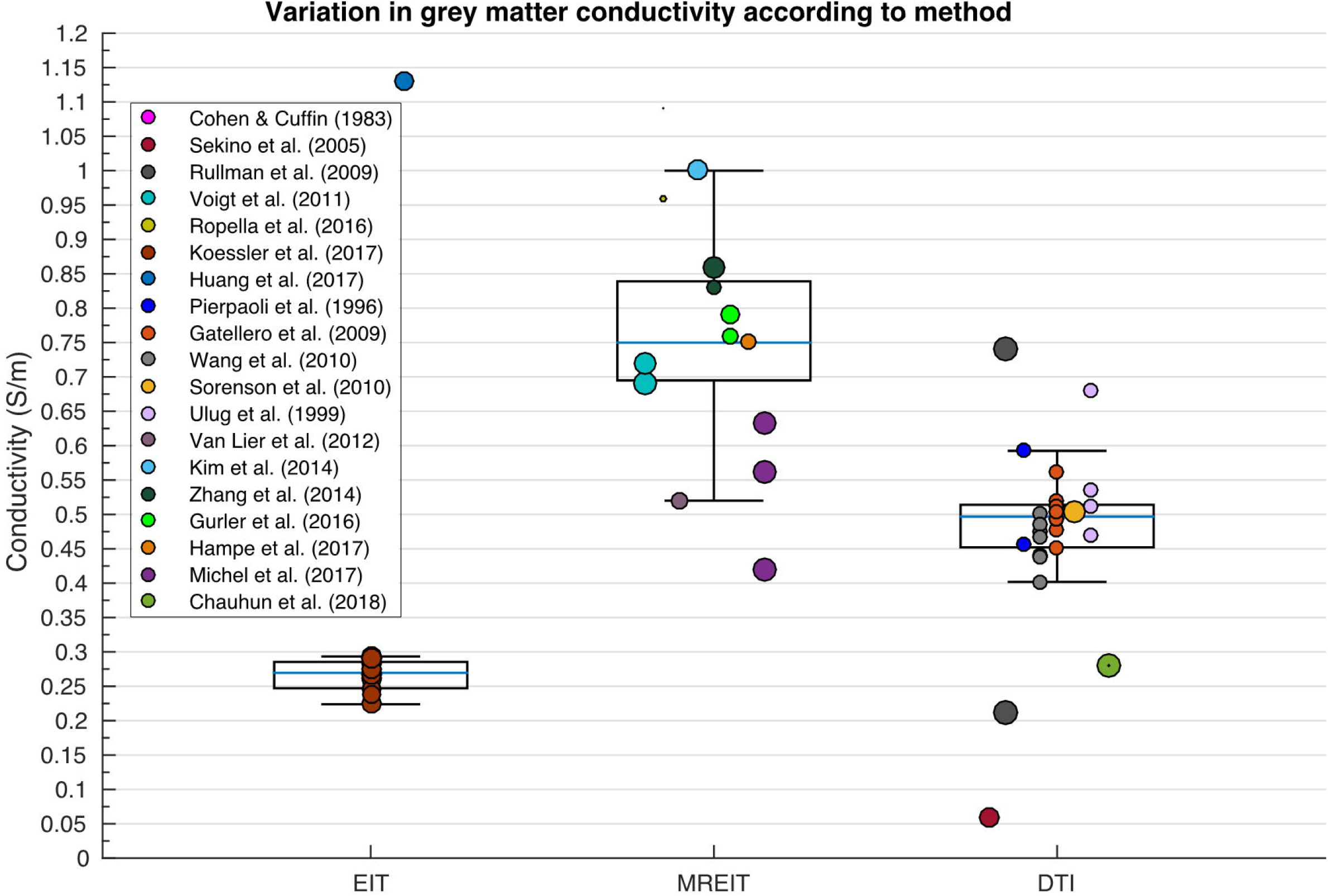
Boxplot displaying variation in GM conductivity depending on method.

#### 3.6.2 White Matter

A weighted multiple regression analysis revealed variation in isotropic WM conductivity was significantly explained by methodology, condition, frequency, pathology and age collectively [R^2^(5, 36) = .696, p <.001], where values varied significantly according to method [Figure 12; F(4,99) = 34.659, p<.001], condition [F(2, 101] = 30.089, p<.001], pathology [F(2, 101) = 34.437, p<.001), temperature [t(102) = 3.877, p<.001] and frequency [(r(104) = −.362, p=.001]. Furthermore, pathology and age collectively explained a significant proportion of variation in WM conductivity measured perpendicularly [R^2^(2, 14) = .459, p =.014] and in parallel [R^2^(2, 14) = .677, p < .001]. Where perpendicular WM values varied according to condition [F(2, 38) = 36.828, p<.001], temperature [t(39) = 1.105, p=.031] and participant age [r(41)=.638, p=.006], whilst parallel WM measurements differed with condition [F(2, 38) = 9.78, p<.001] and age [r(41) = .520, p=.032].

**Figure 11.**
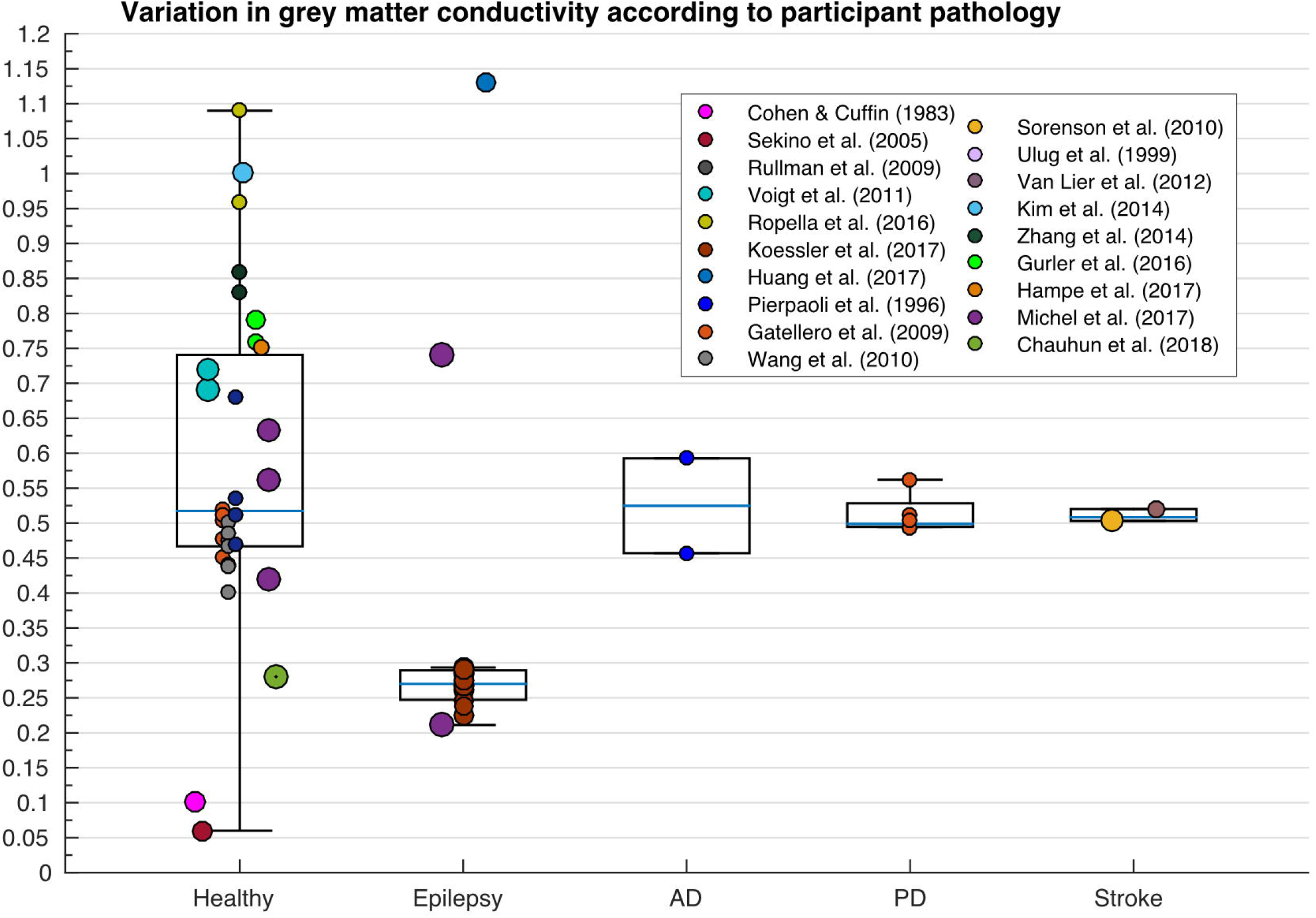
Boxplot displaying variation in GM conductivity depending on pathology (AD; Alzheimer’s Disease; PD; Parkinson’s Disease).

**Figure 12.**
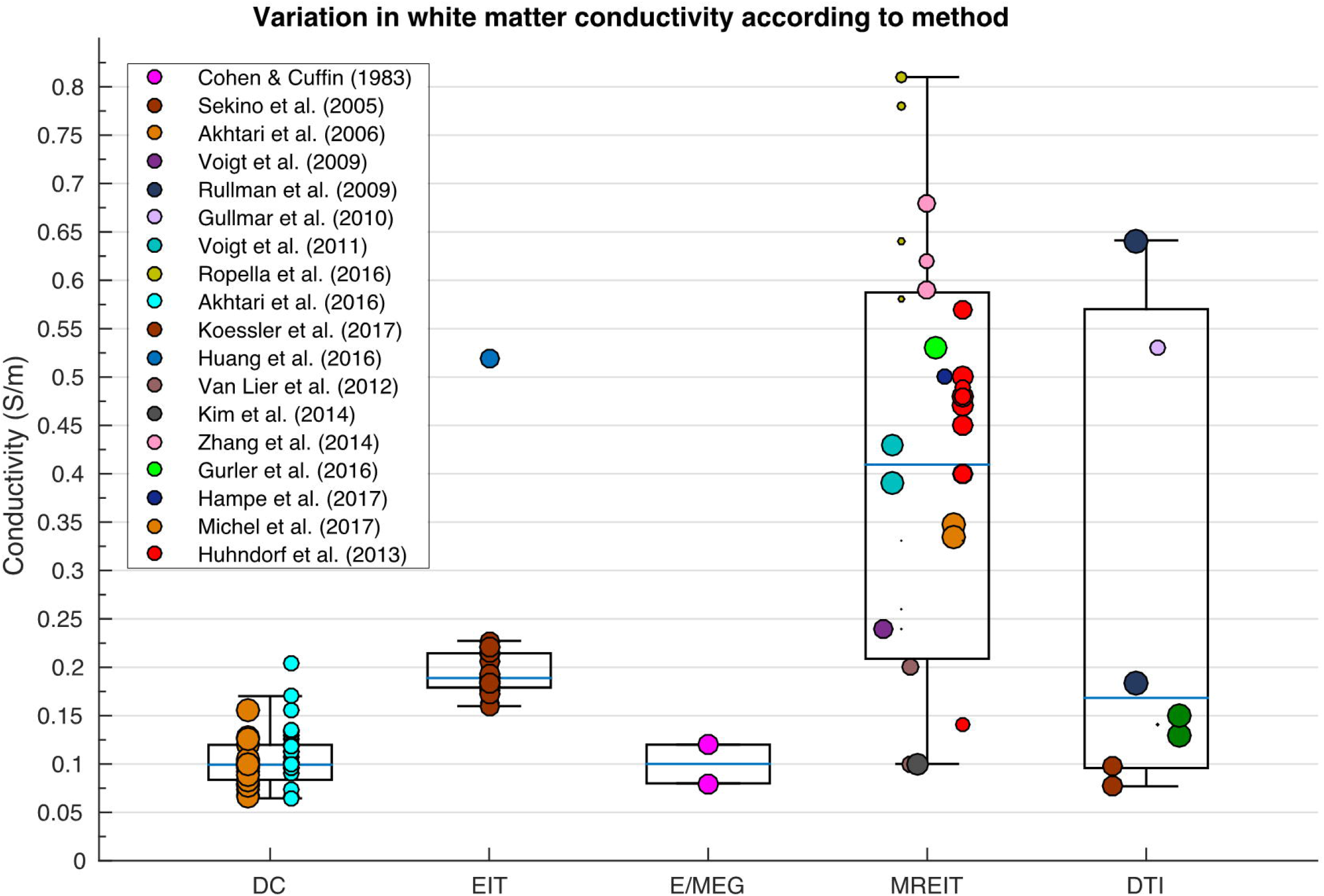
Boxplot displaying variation in WM conductivity according to method.

### 3.7 Brain to skull conductivity ratio (BSCR)

Variation in BSCR calculations was significantly predicted by methodology, frequency, pathology and age collectively, in the weighted regression analysis [R^2^(4, 26) =.302, p =.046]. Figure 13 displays the variation of BSCR according to method, although a comparison of means revealed no significant differences between employed technique. Additionally, BSCR values correlated positively with participant age [Figure 14; r(51) = .376, p = .014].

**Figure 13.**
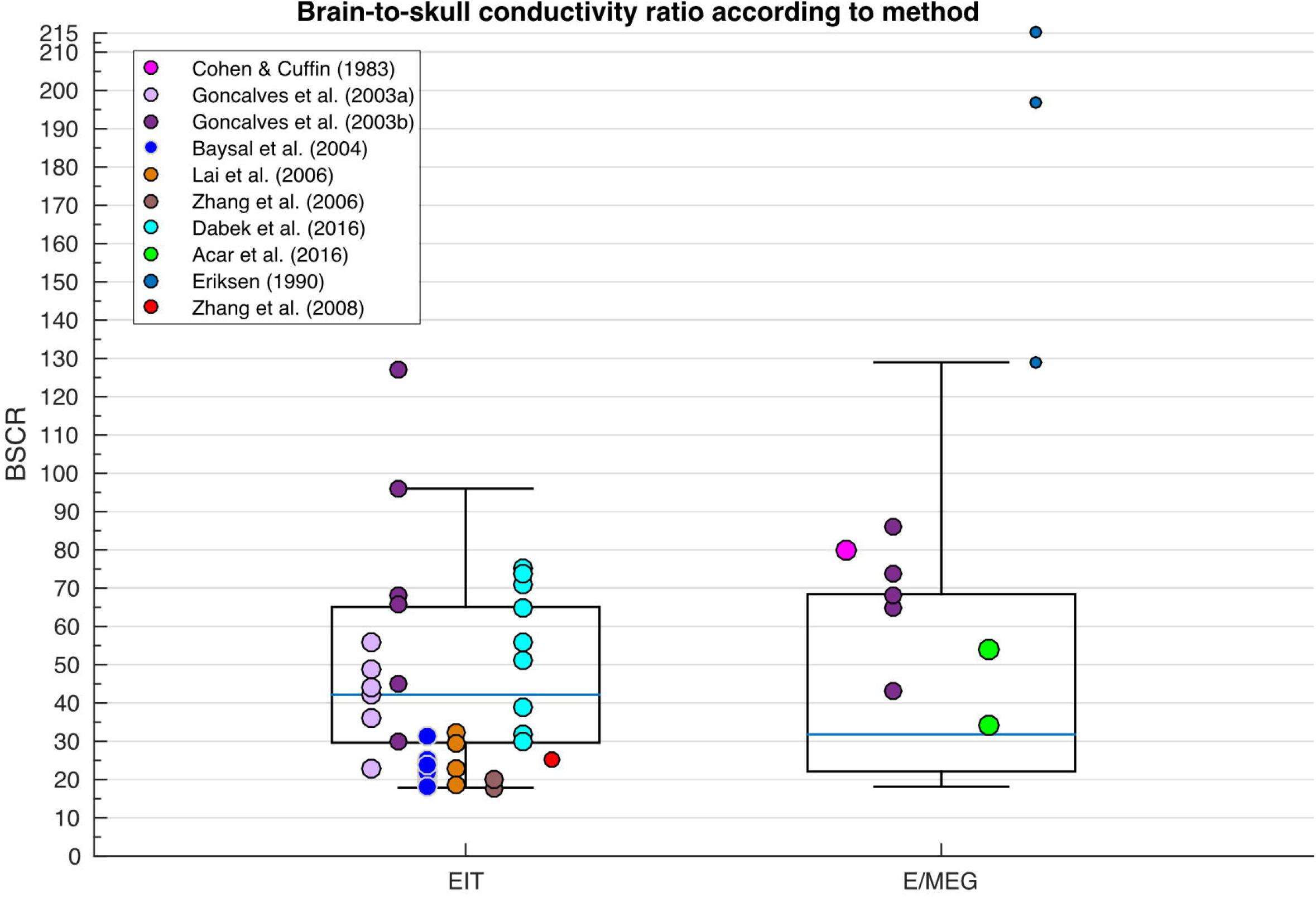
Boxplot displaying variation in BSCR depending on method.

**Figure 14.**
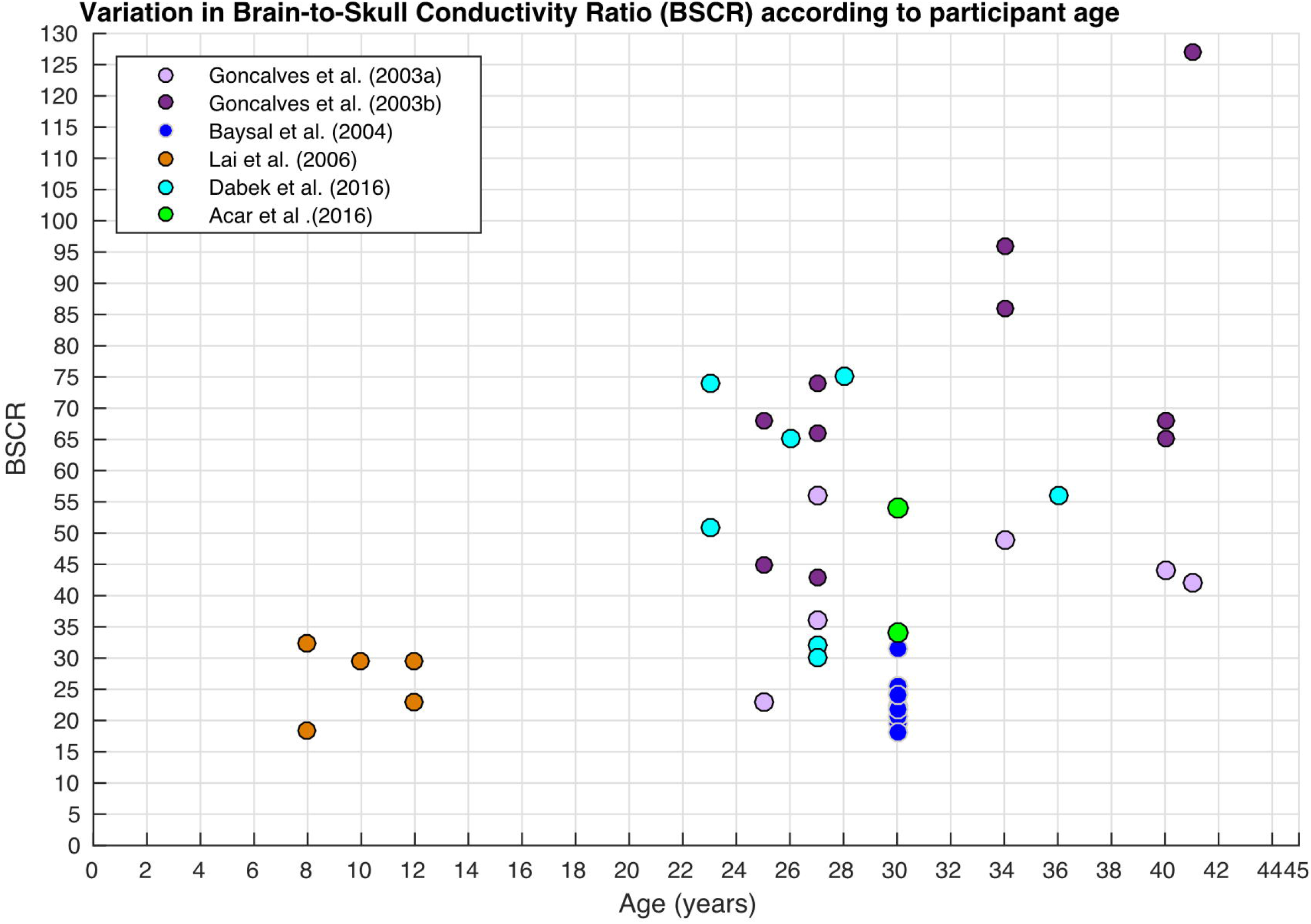
Scatter diagram displaying BSCR as a function of participant’s age.

## 4 DISCUSSION

The current study systematically investigated variation in conductivity of 17 different head tissues as reported in 56 research papers, identified through a literature search of three relevant databases. The mean, standard deviation, minimum and maximum were calculated for each tissue type (Table 3). In addition, we computed the weighted average mean, which provided an optimum (and therefore suggested) value when conductivity is unable to be obtained on an individual basis. The weighted average mean and standard deviation (in S/m) for each tissue type were: scalp = 0.41± 0.18, whole skull = 0.02 ± 0.02, spongiform skull layer = 0.048 ± 0.07, whole compact skull layer = 0.005 ±0.002, outer compact = 0.005 ± 0.003, inner compact = 0.007 ± 0.004, CSF = 1.71 ± 0.3, GM = 0.47 ± 0.24, WM = 0.22 ± 0.17, WM perpendicular = 0.12 ± 0.05,WM parallel = 0.12 ± 0.09, Blood = 0.57 ± 0.11 and BSCR = 50.4 ± 39. The differences between values for each tissue were statistically tested against methodological and participant demographical variables to reveal significant predictors. Inadequate data was available for muscle, fat, blood, the epileptogenic zone, the dura layer, the cerebellum and brain lesions to carry out a multiple regression. Collectively, the independent variables (related to both methodology and demographics) insignificantly explained variation in the scalp, the compact layers of the skull, cerebrospinal fluid, the whole-brain and grey matter. In contrast, variation in whole-skull conductivity could significantly be explained by all the IV’s collectively, where values were revealed to specifically differ significantly depending on method. Variation in the conductivity of the spongiform bone layer of the skull was significantly predicted by condition, frequency, pathology and age, where values were significantly different according to condition and temperature. Despite insignificant results for regression analysis, GM significantly differed depending on method and pathology. Variation in isotropic WM electrical conductivity was significantly predicted by methodology, condition, frequency, pathology and age collectively in the regression model, where values significantly diverged according to method, condition, pathology, temperature and frequency. A significant proportion of variation in WM conductivity measured perpendicularly and in parallel was further explained by pathology and age collectively. Specifically, perpendicular WM values differed with condition, temperature and age, whilst parallel WM measurements fluctuated with condition and age. Lastly, the meta-regression revealed the BSCR could be significantly attributed to variation in methodology, frequency, pathology and age collectively, revealing a positive correlation between the ratio and participant’s age.

### 4.1 Data exclusions

Explanations for the presence of outliers and reasons for any data exclusion in the current meta-analysis are further discussed. Firstly, data acquired at frequencies higher than 1000Hz were removed, as these conditions were deemed unnatural, considering the bandwidth of most neuronal signals is 1Hz - 1KHz, with resonant frequencies of <100Hz for the brain (Groppe *et al.*, 2013) and <1000Hz for the skull (Håkansson *et al.*, 1994). Conductivity results obtained from (Baysal & Haueisen, 2004) employing a conventional least-squares estimator (LSEE) on E/MEG data were further revealed from the weighting algorithm, as stated by the authors as being “unrealistic [negative resistivities] and unstable” (Baysal & Haueisen, 2004). These unstable results were evident from the large standard deviation percentage (>5000% for the scalp, >200% for the skull and >240% for the brain). The authors suggested such inaccuracies occurred from the use of LSEE linearization in a highly non-linear problem, and hence were omitted from their own analysis. Furthermore, skull conductivity values reported by (Hoekema *et al.*, 2003) were elevated approximately tenfold, suggested to be “as expected [due to measurements] in non-physiological circumstances” (namely, saline-coated cadaver), however were not excluded from the current analysis as methodology was in line with previous cadaver studies and therefore should yield similar results. Huang and colleagues (2017) yielded significant outliers for the scalp, skull, whole-brain, GM and WM, where median optimal conductivity obtained by fitting model outputs (from literature) to recordings. These deviations may be explained by the use of an optimisation EIT approach where “best-fit” values were free to compensate for all inaccurate/simplified sources, tissue segmentation errors, changes in electrode location, etc., and therefore cannot reflect “true” conductivity values. Outliers were additionally revealed for the spongiform (but not compact) skull layer by Fernández-Corazza and colleagues (2017), which employed boundary EIT (bEIT) for reconstructing the electrical conductivity for a subset of the regional tissue parameters. The authors acknowledged low sensitivity of bEIT to spongiform variations, due to the relatively small proportion of spongiform to head volume and concluded such approximations may be difficult for unbiased bEIT estimators but remain valid for compact bone estimates. Substantial outliers were further exposed for CSF, GM and WM (as well as subsequent EMA whole-brain calculations) in Ropella & Noll’s (2017) paper, which employed MREIT and revealed standard deviations of 80.2 - 518.8%. The validity and reliability of MREIT as a method is later discussed. The Quality Analysis in the current paper attributing a confidence weighting to each value, is deemed acceptable to consider the large standard deviations leading to outliers. All outliers were therefore included in order to fully account for and explore reasons for variation in values.

### 4.2 Scalp

Additional deviations, although not classified as outliers, were revealed from (Gutierrez *et al.*, 2004) for the scalp, where maximum likelihood estimation (MLE) was used to estimate the electrical conductivity from E/MEG measurements. Such results may be due, in part, to the necessity of accurate source location and head geometry knowledge to avoid estimation bias. The authors suggest the use of a Bayesian approach, which permits incorporating *a priori* information on conductivity distribution to reduce bias. Realistic measurements (excluding outliers and deviations, as discussed) of scalp conductivity, for example, ranged between 0.25 S/m and 0.435 S/m, which could not be attributed to any of the IV’s. Such variation can be relevant in source localisation based on E/MEG. In particular, dipole sources close to measurement electrodes are sensitive to scalp conductivity (Gençer and Acar, 2004; Goncalves *et al.*, 2003). These results coupled with those of the current meta-analysis indicate that assuming scalp conductivity from the literature is not only inaccurate but can lead to E/MEG source mislocalisation. These errors do not appear to be explainable by anything other than individual variability and hence personalised models of scalp conductivity should be considered to improve E/MEG activity localisation.

### 4.3 Skull

According to the meta-regression, variation in whole-skull conductivity can be accounted for by differences in all of the independent variables (methodology, condition, temperature, frequency, pathology and age), with specific differences between methodology and condition. Such significant results, however, may be due to overfitting of the data and meta-regression parameters employed. Future research could utilise machine learning techniques in order to refine the regression analysis and determine the most influential variables. This analysis was beyond the scope of the current paper and ideally would require more data which is not presently available.

Values obtained from excised tissue in non-physiological circumstances after undergoing various processing, may change the electrolyte concentration and therefore skull conductivity (Akhtari *et al.*, 2002). Considering the contrast between a saline-soaked processed cadaver and live skulls that remain in natural conditions between the scalp and meninges, differences in conductivity values post-mortem should be expected. Early research determined conductivity of rat femurs to decrease by a factor of 2.5 - 3 from live-to-50 hours postmortem (Kosterich, Foster, & Pollack, 1983, 1984). This was further corroborated in the human skull, indicating a scaling factor of 2.5 – 4 from live-to-post-mortem (Wendel *et al.*, 2006). These results are validated by Rush and Driscoll’s mathematical model stating the skull conductivities dependency on the fluid (i.e. saline vs naturally positioned between meninges, surrounded by blood/CSF, etc.) permeating it (Rush & Driscoll, 1969), suggesting, a live skull to have higher conductivity than a saline-soaked (with conductivity 1S/m) cadaver skull. It is consequently unsurprising *in vitro* values of skull conductivity differ from *in vivo* values – predicting increased conductivity *in vivo* (Akhtari *et al.*, 2000; Law, 1993). The results from the previous literature coupled with the current meta-analysis, therefore indicate skull conductivity should be measured in *in vivo* (i.e. at body temperature, within the live head, etc.) in order to avoid bias and increase reliability.

Within *in vivo* reports, despite all under similar conditions, skull conductivity values obtained using E/MEG appeared to be elevated compared to those employing EIT. These results may be explained by the use of statistically constrained estimating algorithms for E/MEG measurements, for which conductivity is optimally estimated from E/MEG arrays acquired during electrical nerve stimulation. Furthermore, whole-skull conductivity was found to vary as a function of frequency, although the nature of the relationship was unclear. Previous literature has examined skull conductivity in wider frequencies, revealing a positive relationship, especially at frequencies higher than 10KHz (Gabriel, Gabriel, & Corthout, 1996b; Tang *et al.*, 2008), and suggest skull conductivity may exponentially increase at high frequencies. Within the relevant range for brain activity, skull conductivity increased by ~6.7% from 11-127Hz (Dabek *et al.*, 2016) and ~13% from 10-90Hz (Akhtari *et al.*, 2002), of which the authors developed a non-linear model for frequency dependence of different skull layers (Akhtari *et al.*, 2003). Variation as a function of frequency may be expected due to interactions between mobile electrolytes (i.e. sodium and chloride) and relatively immobile molecules (i.e. proteins and blood components) which affect the relaxation rate of conductivity dependent on the currents frequency (khtari *et al.*, 2002; Latif *et al.*, (2010). Importantly, this frequency dependence has been implicated in causing the volume conductor to act as a low pass filter, which potentially contributes to E/MEG forward solutions errors and may therefore reduce accuracy in E/MEG source localisation.

Furthermore, results exploring differences according to age in the BSCR (see Figure 14) have implications for skull conductivity variation. Research has indicated skull, rather than brain, conductivity plays a larger role in BSCR values (Goncalves *et al.*, 2003), consequently suggesting skull conductivity varies with age. These results are discussed further in section 4.3.1 (layered skull conductivity) and 4.6 (BSCR).

#### 4.3.1 Layered skull

The majority of previous studies simplify the skull as a homogeneous layer, not accounting for differences in conductivity between the compact (upper, lower) and spongiform layers of the skull. Distinct conductivities for the three layers of the skull has been previously indicated (Akhtari *et al.*, 2000; Akhtari *et al.*, 2002; Fernández-Corazza *et al.*, 2017; Tang *et al.*, 2008), as supported by the current meta-analysis (see figure 2). This is unsurprising considering the higher prevalence of fluid filled pores and cavities in spongiform (and hence higher conductivity) compared to compact bone. Importantly, neglecting inhomogeneous estimations for a tri-layer skull has yielded significant errors in source localisation irrespective of model parameters (Dannhauer *et al.*, 2011; Haueisen *et al.*, 1999; Haueisen *et al.*, 2002; Ollikaineet *et al.*, 1999; Pohlmeier *et al.*, 1997). These authors have thus concluded realistic modelling of the skull layers to be necessary for accurate E/MEG source localisation. The current meta-analysis, however, revealed that variations exist between individuals even whilst considering a tri-layer skull. Variation in the electrical conductivity of the spongiform skull layer was revealed to be significant and attributed to deviations in condition, temperature, frequency, pathology and age. Suggesting true values for conductivity of the spongiform layer will not only depend on methodological parameters but also individual demographics. Likewise, although the compact layers of the skull were insignificantly predicted by any of the parameters, large variation was still evident. This further elucidates the hypothesis that conductivity values fluctuate between individuals and support the suggestion for personalised models of skull conductivity.

Interestingly, a relationship with age was to be expected, regardless of homogeneity, considering the presence of the fontanels and open sutures (un-ossified bone filled with cartilage, chondroid and vascular tissue) which may remain unfused for several years (Hansman, 1966). Firstly, bone formation (remodelling) from direct laying down or cartilage, known as ossification, remains approximately 50% complete at birth, not reaching 100% until age 20. This indicates a higher proportion of cartilage and “soft” (trabecular) bone, consisting of higher water and less lipid content (Silau, Fischer, & Kjaer, 1995), in the neonatal skull, which linearly decreases until fully ossified (Christie, 1949). Furthermore, neonatal brain growth occurs at a rate higher than bone ossification, resulting in four fontanelles between the skull plates. The fontanelles, although relatively small at birth, become larger within the first few months (up to 3 cm) until closing at approximately 18 months old (Hansman, 1966). As well as inter-subject variability, the size and width of fontanelles significantly vary in children with central nervous system pathology. Moreover, unclosed sutures in an infant skull, although present in an adult skull, are wider near the fontanelles, and may not close for several years in healthy children (Erasmie & Ringertz, 1976). In addition to the fontanelles, the sutures close at various times of development. The frontal suture, for example, normally fuses between 3-9 months old (Vu *et al.*, 2001), whereas the sphenofrontal suture (lying between the sphenoid bone and posterior horizontal orbital plates) and sagittal suture (connecting the two parietal bones) typically close by age 15 and 22 years, respectively. Moreover, the squamosal sutures (connecting the temporal squama and parietal bone), do not fully close until 60 years of age (Vijay Kumar *et al.*, 2012). Lastly, the skull thickness increases with age, from 2-3mm at birth to 3-6mm during early adulthood (Despotovic *et al.*, 2013; Hansman, 1966). This increase however is non-linear, slowing down towards 3 years of age, and is non-uniformly distributed throughout the skull, with higher thickness in occipital than frontal and parietal regions (Li *et al.*, 2015).

Taken together, the structural differences between the neonatal and adult skull elucidate differences in skull conductivity to be expected. As such, previous studies have revealed higher skull conductivity for infants compared to adults, (Gibson, Bayford, & Holder, 2000; Pant *et al.*, 2011) and an inverse correlation between skull conductivity and thickness with increasing age (Gibson *et al.*, 2000). The inner and outer compact layers are thought to become thicker, whilst the spongiform layer becomes thinner with age, hence decreasing whole-skull conductivity. Additionally, paediatric skull tissue ordinarily contains greater quantities of ions and water, compared to calcified cranial bones of adults, hence higher conductivity may be expected (Schönborn, Burkhardt, & Kuster, 1998). Further support from animal studies have revealed skull conductivity to decrease with age, for example, in the rat (Peyman, Rezazadeh, & Gabriel, 2001), pig (Peyman *et al.*, 2007) and cow (Schmid & Überbacher, 2005). Although this expectation was not confirmed in the current meta-analysis, for accurate source localisation, skull conductivity variations with age should be taken into consideration. For example, Lew and colleagues assessed the effect of sutures and fontanels on E/MEG source analysis, where omission produced a maximum position error of 3.6mm and 0.6mm for tangentially- and radially-oriented sources, respectively (Lew *et al.*, 2013). Elevated EEG localisation errors with respect to MEG were further replicated in simulation studies (Flemming *et al.*, 2005), suggesting an advantage in employing MEG for developmental studies. The literature however suggests both modalities require individualised, or in the least an infant-specific volume conductor model to accommodate for relevant developmental changes (Bystron, Blakemore, & Rakic, 2008; Rakic, 2006; Song *et al.*, 2013).

Following from this, development of the human skull does not cease after infancy, but continues to undergo remodelling, microstructural, density and histological changes until death, further impacting conductivity. Firstly, total cranial thickness has been observed to increase with age (Todd, 1924) notably related to increase in diploë thickness (Hatipoglu *et al.*, 2008; Sabancioğullan *et al.*, 2012), which in one study was accompanied with inner and outer compact thinning (Skrzat *et al.*, 2004). An increase in diploë (and hence spongiform bone) thickness would suggest conductivity of the skull to increase with age, as revealed by (Tang *et al.*, 2008). Recently, Antonakakis and colleagues (2018) revealed a trend between participant age and skull conductivity but noted the small sample size (n=15) and large inter-subject variability rendered robust conclusions difficult and inadequate. Further results have also been inconsistent, finding no such relationship between skull thickness and age (Ishida & Dodo, 1990; Lynnerup, 2001; Lynnerup, Astrup, & Sejrsen, 2005; Pensler & McCarthy, 1985; Sullivan & Smith, 1989). The presence of suture lines, not limited to infants, was furthermore shown to increase conductivity of the skull sample, by providing a path of high conductance (Law, 1993; Tang *et al.*, 2008). Additionally, the percentage of spongiform bone within the skull was positively correlated with skull conductivity (Tang *et al.*, 2008), whilst, skull thickness, which is non-uniform within and between individuals (Lynnerup, 2001; Lynnerup *et al.*, 2005), was inversely correlated with scalp potentials (Chauveau *et al.*, 2004). One paper, for example, revealed that a 20% and 40% decrease in skull thickness resulted in a 5-10% and 20-25% decrease in conductivity, respectively (Lai *et al.*, 2005). Insufficient results were available to analyse the influence of sutures or skull thickness in the current review; however, the discussed structural deviations further illuminate the importance of employing individualised models of head conductivity.

The influence of skull conductivity and segmentation inaccuracies has been explored extensively, revealing overwhelming source localisation errors for EEG (Lanfer *et al.*, 2012; Montes-Restrepo *et al.*, 2014; Wolters *et al.*, 2006) and MEG (Cho et al., 2015; Lau *et al.*, 2016) of up to 2cm. E/MEG source analysis can importantly be utilised during presurgical epilepsy diagnoses, together with MRI data. Aydin and colleagues (2017) recently developed a multimodal technique, of which they emphasised the importance of individualised high-resolution finite element head models with WM anisotropy modelled from DTI and individually calibrated skull conductivity, alongside combined E/MEG and MRI information. Importantly, they note creating such realistic head models may not always be feasible, and therefore recommend, at minimum, skull conductivity to be individually adjusted to improve combined E/MEG source analysis. Variations in skull conductivity have been found to impact transcranial electric stimulation focality and dose (Santos *et al.*, 2016; Schmidt *et al.*, 2015; Wenger *et al.*, 2015), with one study revealing an error of 8% in dose (Fernández-Corazza *et al.*, 2017). Such inaccuracies are clinically relevant, particularly regarding source estimation for refractory epilepsy (Brodbeck *et al.*, 2011) and determining electrical current dose required for treatment of epilepsy (Berényi *et al.*, 2012; Liebetanz *et al.*, 2006), depression (Kalu *et al.*, 2012) and other psychiatric disorders (Brunoni *et al.*, 2013). Of note, Dannhauer and colleagues (2011) investigated variations in layered skull structures for EEG forward modelling and revealed an inhomogenous but not isotropic modelling to be of most importance. For optimum skull modelling, they recommended assigning each skull voxel to a tissue type (compact or spongiform bone) with individually estimated or measured conductivity values. To optimise source localisation accuracy, we therefore recommend employing personalised models of skull conductivity to sufficiently consider individual variability.

### 4.4 CSF

The results of the current meta-analysis are in line with previous report from Baumann and colleagues (1997), displaying significant variation in CSF conductivity dependent on temperature. They revealed 23% higher conductivity at body (37°C), approximately 1.79 S/m, than room (25°C) temperature, which corresponded to the temperature coefficient of 2% per 0.1ml of potassium chloride (comparable to CSF conductivity and ion concentration (Fishman, 1992; McGale *et al.*, 1977; Wu, Koch, & Pratt, 1991). The result from Baumann *et al*.’s (1997) study is frequently considered as a reference value for CSF conductivity. However, the current meta-analysis has revealed large variation in measurements, potentially suggesting instability in CSF conductivity between individuals. The majority of such variation stems from values obtained using MREIT. One such study, (Ropella & Noll, 2017) reported values with standard deviations up to 518% of the mean, which indicates instability in the methodology. The combination of large standard deviations within and between studies potentially call into question the validity of MREIT for measuring conductivity of the human head – this is further discussed in the grey matter subsection (4.5.1). Further deviating from Baumann’s approximation (1.79 S/m), (Cohen & Cuffin, 1983b) reported a considerably lower conductivity for CSF (1.39 S/m). These results may be explained by their use of optimum estimation, rather than direct measurements; where conductivity was adjusted so the maximum potential in a theoretical EEG map matched the experimental equivalent. Considering the variation due to methodological error, CSF appears to be relatively stable between individuals, with an average conductivity converging around Baumann’s results. Despite this, figure 8 suggests CSF conductivity ranges between ~1.5 – 2 S/m, elucidating the need for individualised measurements. Deviation in CSF conductivity in that range however, may not significantly affect E/MEG forward and inverse modelling solutions. Future work could therefore employ a sensitivity analysis to explore the influence CSF conductivity has on electromagnetic source localisation and determine the necessity of individualised models.

### 4.5 Whole-Brain

The meta-regression failed to explain variation in conductivity of the brain as a homogeneous layer, however values were revealed to significantly vary according to the employed method. Despite these significant results, variation between acquisition techniques are minimal, where large variations within each method remain evident. These variations further support the suggestion that individual values of conductivity should be obtained. However, assuming homogeneous conductivity over the whole brain is generally considered a vast oversimplification and highly inaccurate. Such an assumption fails to consider differences between GM and WM conductivity, as well as structural variation of GM/WM proportion in the brain. Early research determined GM to contain higher proportions of water than WM (Stewart-Wallace, 1939), demonstrating expected higher electrical conductivity for GM compared to WM. Additionally, extensive literature has shown GM and WM volume to vary with development (Giorgio *et al.*, 2010; Groeschel *et al.*, 2010; Miller, Alston, & Corsellis, 1980) and pathology; i.e. multiple sclerosis (Sastre-Garriga *et al.*, 2005), Alzheimer’s Disease (Salat, Kaye, & Janowsky, 1999), schizophrenia (Douaud *et al.*, 2007), attention deficit-hyperactivity disorder (McAlonan *et al.*, 2007), among others. These observations further support the use of individualised models of head volume and conductivity profiles for the most accurate representation of the human head.

#### 4.5.1 Grey Matter

The questionable validity of MREIT for estimating conductivity was further reinforced considering the number of outliers (Ropella & Noll, 2017), coupled with elevated values (Voigt *et al.*, 2011) for GM conductivity acquired with MREIT. Measurements utilising MREIT were over twice those from EIT and approximately one and a half times those of DTI values. Significant variation in GM conductivity may be further explained by increased DTI values relative to EIT. Firstly, the Tuch *et al.* (1999) model derived the conductivity tensor from the water diffusion tensor through differential effective medium approximation (EMA), which uses an electromagnetic depolarisation factor to consider the impact of cell geometry to overall conductivity. This depolarisation factor was originally developed for WM structure, consisting of myelinated pyramidal cells, and therefore may not be completely translational to GM. Tuch and colleagues’ later paper (2001), however, utilised the EMA method to show there exists a strong linear relationship between the conductivity and diffusion tensors, regardless of tissue type. They generated a conductivity tensor image, where conductivity could be assigned to three groups: GM, WM parallel or WM perpendicular to the fibre tract. Their results indicate the EMA method is appropriate for GM conductivity estimation, despite having lower anisotropy than WM. The established linear relationship mapping the diffusion to electrical conductivity tensor employed (Tuch *et al.*, 1999; Tuch *et al.*, 2001) has been further validated in a silk yarn phantom (Oh *et al.*, 2006). However, (Rullmann *et al.*, 2009) detected the use of Tuch’s scaling factor would have generated values 3.5 times greater than isotropic values (taken from (Ramon *et al.*, 2006)). For this reason, they chose to employ a volume constraint approach with scaling factor 0.21 (compared to 0.844) which minimised differences between isotropic and anisotropic EEG forward modelling in their study. Similarly, (Sekino *et al.*, 2005) estimated the effective GM conductivity from only the fast diffusion component (attributed to extracellular fluid), rather than both the fast and slow (attributed to intracellular fluid) components. The produced conductivity maps were therefore not simply linearly scaled diffusion maps, and hence may explain their considerable low GM conductivity measurement (0.06 S/m). Importantly, the deviations in GM conductivity dependent on the chosen diffusion tensor method are acknowledged, emphasising the non-trivial nature of relating the diffusion and electrical conductivity tensors. Future studies should examine this relationship, in order to accurately determine a realistic scaling factor to improve conductivity tensor estimations.

Additionally, Rullmann and colleagues (2009) results were limited to one paediatric participant with epilepsy, of which his age may have influenced the increased brain conductivity. Higher conductivities in paediatric brains are perhaps expected due to the general abundance of water in GM (Dobbing & Sands, 1973). The current meta-analysis failed to find a significant correlation between GM conductivity and age, however, considering normal GM development and the frequently observed decrease in mean diffusivity of GM with age (Pal *et al.*, 2011), further research may expose such a relationship.

In line with this observation, participant pathology significantly affected GM conductivity, but was not a significant predictor in the regression model. Large variation can be seen within and between different participant pathologies (Figure 11), but no clear conclusion could be made. This is perhaps due to the limited number of values available for each classified pathology, reducing the statistical power. In tumours, for example, firstly, all values may not have been made from healthy tissues (due to the diffusing nature of malignant tumours), but also an increased conductivity in “healthy” GM tissue may also be a consequence of tumour cysts increasing the water/CSF concentration in nearby tissues. Previous literature has indicated abnormalities in GM volume, structure, myelination and topography in Multiple Sclerosis, Parkinson’s Disease, Alzheimer’s Disease and other forms of dementia (Compta *et al.*, 2012; Frisoni *et al.*, 2007; Geurts & Barkhof, 2008), as well as abnormalities in psychiatric and developmental disorders (Greimel *et al.*, 2013; Job *et al.*, 2005; Wise *et al.*, 2017). It is therefore unsurprising that GM conductivity varied with participant pathology. Due to the unknown nature of this variation with disease and age, the use of individualised models of head conductivity that are inhomogeneous and anisotropic are especially essential for electromagnetic source imaging (Birot *et al.*, 2014). Increasing the feasibility and accessibility of this could be explored with further research involving DTI parameters.

#### 4.5.2 White Matter

The current meta-analysis failed to explain variation in anisotropic WM conductivity, however this may be due to the limited sample size available for analysis. The crucial consideration of anisotropic conductivities has been more recently determined, where neglecting WM anisotropy produced EEG-localisation errors of ~1.6 - 5.1mm and 4.72 - 8.8mm for radially- and tangentially-oriented sources, respectively (Anwander *et al.*, 2002; Gullmar *et al.*, 2010), with one study reporting a maximum error of 26.3mm (Hallez *et al.*, 2005). Additionally, disregarding anisotropy had a large influence on the induced electric fields from TMS (De Lucia *et al.*, 2007), whilst WM anisotropy influenced the electrical potential distribution following application of deep brain stimulation (Butson *et al.*, 2007; McIntyre *et al.*, 2004). Uncertainty in WM conductivity also had a significantly large effect on tDCS stimulation amplitudes and current density estimations, which were especially pronounced in the auditory cortex, implying orientation to be a determining factor in tDCS applications (Schmidt *et al.*, 2015).

Isotropic WM conductivities however were found to vary dependent on method, measurement condition, pathology and age, additionally diverging with temperature and frequency. As previously observed, MREIT produced elevated WM conductivity values with larger variation between values compared to other methods, supporting the uncertainty and apparently low reliability of MREIT for estimating conductivity. Furthermore, DTI values produced similar, largely varying results. As previously discussed by Rullmann and colleagues (2009), elevated conductivity values from DTI may be a result of an overestimated scaling factor from the diffusion tensor to the conductivity tensor, proposed by the authors. In line with this, Akhtari and colleagues (2006) failed to verify this relationship and instead revealed an inverse linear relationship with a scaling factor of −0.367, and considerable variability between values. For this reason, Gullmar and others (2010) compensated for isotropic variation by normalising the conductivity tensors (by calculating the anisotropy ratio between eigenvalues) in one model and using fixed artificial anisotropy ratios to preserve diffusion tensor orientation, in another model (volume constraint model). By comparing both models with Tuch’s, they revealed that employing the latter significantly affected MEG and EEG forward computations differently by changing the mean scalar representation of isotropic tensors. This study, alongside previous investigations, emphasised the importance of modelling anisotropy, but has also insinuated further, more detailed research should explore the linear relationship between the diffusion and electrical conductivity tensors.

Additionally, differences were revealed between perpendicularly- and parallelly-oriented WM conductivities that are due to the results being obtained from different papers. Of note, the minimum WM_par conductivity value (0.0543 S/m) is less than the minimum WM-perp value (0.0620 S/m), with only small differences reported between the means. These results are not indicative of WM conductivity themselves, but instead highlight the variation in values between studies. Moreover, this may reflect the differences between the methods employed for approximating the conductivity tensor from the diffusion tensor, due to the fact that WM is more anisotropic than these results indicate. A recent paper (Wu *et al.*, 2018) reviewed the current anisotropic conductivity models of WM based on DTI. The linear relation model (i.e. Tuch’s model) was discussed as not directly considering the impact of geometrical brain tissue structure, whilst the Wang-constraint (Wang *et al.*, 2008) and volume-constraint models, both of which assume diffusion and conductivity tensors share the same eigenvalues (similar to Tuch’s model), ignore brain tissue heterogeneity but can relate anisotropy and physiological structure. The equilibrium model (Sekino *et al.*, 2005), which decomposes extracellular and intracellular diffusion, can be less accurate as extracellular diffusion may result difficult to quantify (Jones *et al.*, 2018). Wu *et al.* (2018) determined that obtaining the conversion coefficient between the anisotropic conductivity tensor and the diffusion tensor eigenvalues to be of most importance. Further, they concluded the optimum model to be the electrochemical model, which calculates the conversion coefficient according to the concentration of charged particles in interstitial fluid and has the added benefit of being able to calculate a conversion coefficient for GM and avoids having to consider the effect WM structure has on water molecules and electrical charges. They noted however, that the models are not contradictory, but instead complement each other to inform the relationship between conductivity and diffusion tensors. This emphasises the need for more research to elucidate the prime conversion coefficient before robust conclusions can be made. Exploration and evaluation of different diffusion-to-conductivity methods are beyond the scope of the current meta-analysis, however, considering the variable results in combination with Wu *et al.* (2018) analysis, caution should be taken when applying conversion algorithms between conductivity and diffusion tensors.

Furthermore, as formerly discussed, measurements obtained *in vitro* or *ex vivo*, as well as at room temperature, are likely to differ from *in vivo* results due to the non-physiological conditions. It is therefore recommended investigations should be completed at body temperature, *in vivo* and at frequencies in line with resonant frequencies of the brain (<100Hz). Subsequently, a significant correlation between WM conductivity and applied frequency may have only be revealed due to values obtained at 500Hz. A large pool of conductivity values were obtained at 500Hz, whilst the remainder of values were measured at frequencies <150Hz, hence, the higher conductivity values at a considerably higher frequency than all other results may have skewed the data to reveal a positive relationship, which may not otherwise be there. Additional conductivity values measured with frequencies between 150Hz and 500Hz are needed to further elucidate the presence or not of a significant relationship.

WM conductivity values were additionally revealed to vary with pathological condition, however insufficient results were available to extract clear conclusions. Such variation with pathology is nevertheless, expected, for example intracranial pathology from epilepsy patients potentially alters cytoarchitecture of affected and non-affected areas, such as cortical neuronal disorganisation and surplus WM cells (Mathern *et al.*, 1999). Akhtari and colleagues (2006) suggested pathological changes in myelin and diseased-active cells may disrupt cell geometry organisation, which, when coupled with histological demyelination and cell population alterations, may increase proton diffusion as it is no longer constrained by myelin walls or tight organisation. Interestingly, a significant relationship between fluctuations in DTI eigenvalues and histological alterations in temporal lobe epilepsy has been found (Kimiwada *et al.*, 2006). Furthermore, extensive research has revealed marked differences in WM structure in numerous diseases and pathologies, such as Multiple Sclerosis, Alzheimer’s Disease, Schizophrenia and Parkinson’s Disease (Bozzali *et al.*, 2002; Burton, McKeith, Burn, Firbank, & O’Brien, 2006; Kubicki *et al.*, 2005; Kutzelnigg *et al.*, 2005). Conductivity and anisotropy surrounding diseased tissues is subsequently likely to affect the generated electrical and magnetic fields from a source, compared to such distributions in healthy individuals (Park *et al.*, 2002; Youn, Park, Kim, Kim, & Kwon, 2003).

Although no significant correlation was revealed, WM conductivity as a function of age was revealed to contribute to the meta-regression model. This observation is unsurprising considering the well-researched nature of WM development with age, suggesting a decline in WM integrity, WM volume, myelination, diffusivity, etc. (Gunning □ Dixon, Brickman, Cheng, & Alexopoulos, 2009; Guttmann *et al.*, 1998; Salat *et al.*, 1999; Schmithorst, Wilke, Dardzinski, & Holland, 2002). The degeneration of WM with age allows for higher CSF and liquid concentrations within the WM, therefore increasing conductivity of the tissue. Considering the large variation of WM with methodology, pathology and participant demographics, assuming conductivity of WM from the literature is clearly insufficient for accurate head conductivity profiles.

Combining the current results for WM and GM conductivity values, a clear discrepancy exists between both tissues, indicating the heterogeneity of the brain. Assuming the brain to have homogeneous and isotropic conductivity, as in research considering the brain as a whole, is therefore insufficient for accurate conductivity profiles. Assuming the brain as a homogeneous conductor consequently results in considerable EEG and MEG source localisation errors (Acar *et al.*, 2016; Awada *et al.*, 1998; Cohen & Cuffin, 1983b).

### 4.6 Brain-to-skull Conductivity Ratio

The ratio between brain and skull conductivity was significantly different for epilepsy compared to healthy participants, however, all BSCR values with epileptic pathology were obtained from paediatric samples. In accordance with this observation, BSCR was revealed to increase with age, suggesting paediatric samples, and hence epilepsy in the current review, to have lower conductivity ratios. In support of previous literature examining the influence of age on skull conductivity, such a relationship is expected, considering increased conductivity and decreased thickness of paediatric skulls. In contrast, paediatric brain tissue contains relatively higher water content and lower myelin deposition than adults (Knickmeyer *et al.*, 2008; Peterson *et al.*, 2003), indicating higher conductivity. Together, higher conductivity of both the skull and the brain in paediatric samples would suggest the brain-to-skull ratio to remain relatively stable throughout age. However, this would require an equal rate of decline for both tissues, which is unlikely the case. The role of skull conductivity for BSCR calculations was elucidated by (Goncalves *et al.*, 2003), who concluded their BSCR discrepancies to be a consequence of skull, as opposed to brain, conductivity variation. Their results indicated brain conductivity to be of less importance when calculating the brain-to-skull conductivity ratio, rendering decline in brain conductivity, whether equal or not to skull conductivity decline, irrelevant.

In contrast, assuming isotropic and homogeneous properties of the skull may contribute to variations in brain-to-skull estimations. For example, current injection at different locations may result in differing impedance distribution within the skull, hence altering BSCR estimations when isotropic compared to anisotropic skull models are used (van den Broek, Reinders, Donderwinkel, & Peters, 1998). However, it is noted that more recent papers acknowledge the importance of segmenting the skull layers, hence the use of skull anisotropy to optimise estimates of layered skull conductivity may not be required. Additionally, the heterogeneity in the skull will evidently introduce conductivity variation when homogeneous models are used, hence contributing to variation in BSCR values. These observations similarly apply for brain homogeneity and anisotropy, which would consequently affect BSCR estimations and contribute to variation. Considering BSCR estimations are clearly dependent on accurate conductivity of the skull and brain, which are subject to large variability, it is suggested personalised models of whole-head conductivity are essential to accurately determine the brain-to-skull conductivity ratio.

## 5 CONCLUSION

The current meta-analysis systematically investigated variation in reported human head electrical conductivity values for 17 different tissue types to evidence deviation within and between tissues, determine any influential factors and evaluate the impact on E/MEG source reconstruction. Adhering to the hypothesis, conductivity was revealed to significantly vary throughout the literature, specifically for the scalp, different layers of the skull, the whole-brain, grey matter, white matter and the brain-to-skull conductivity ratio. To decrease variation and increase stability of conductivity estimates, values should be obtained at body temperature, at frequencies less than 100Hz and in natural, *in vivo* conditions. Conductivity was significantly discovered to vary dependent on participant pathology, suggesting separate values should be acquired for different pathologies. However, further research was suggested to enhance the understanding of pathological effects on the electrical conductivity field. Additionally, electrical conductivity significantly correlated with participant’s age for the skull, WM and BSCR. Previous literature was presented indicating large EEG and MEG source localisation errors occurring as a result of inaccurate conductivity values, particularly when neglecting anisotropy and heterogeneity of the head. The impact of incorrect conductivity on E/MEG forward and inverse solutions, coupled with the high variability of conductivity for each tissue type, suggests that assuming conductivity from the literature is insufficient. In conclusion, to optimise source reconstruction in EEG and MEG, and reduce localisation errors, personalised models of head electrical conductivity should be obtained for each individual.

## Acknowledgements

Knowledge Economy Skills Scholarships (KESS) is a pan-Wales higher level skills initiative led by Bangor University on behalf of the HE sector in Wales. It is part funded by the Welsh Government’s European Social Fund (ESF) convergence programme for West Wales and the Valleys.

## APPENDIX A: PRISMA FLOW DIAGRAM

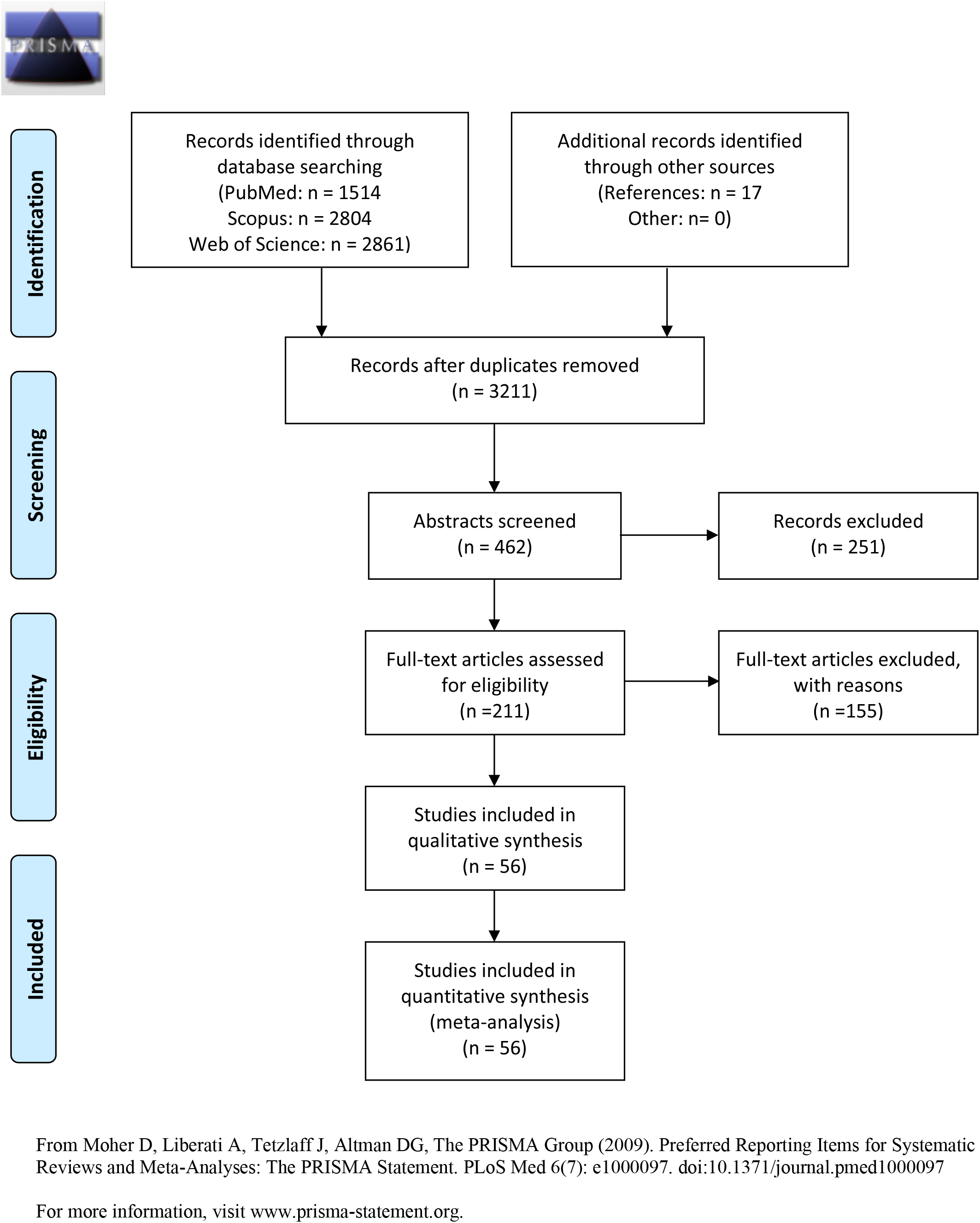

## APPENDIX B: KEYWORDS FOR LITERATURE SEARCH

The inclusion criteria considered Articles, Books, Book Chapters, Corrections, Data Paper, Early Access or Reprints presented or translatable to English. The keyword search was conducted for titles only. Keywords included any variation and combination of “conductivity” (i.e. resistance, impedance, dielectric, electric/current field, electric properties) AND “head tissue” (i.e. head, brain, scalp, skull, cerebral, CSF, dura, white matter, grey matter, brain-skull, brain-scalp, BSCR, lesion). To reduce the amount of retrieved papers, those including unrelated keywords in their titles (i.e. insulin, diabetes, drug, DNA, blight, ship, sea, flower, Kawasaki, train) and non-human animals (i.e. rat, pig sheep, cow, swine, mice, mouse) were excluded from the keyword search.

## APPENDIX C: QUALITY ASSESSMENT PROTOCOL FOR ALL STUDIES

| **Item 1: Were participants appropriately recruited and described?** |  |
| --- | --- |
| Pre-specified protocol | No participant demographics or pathology were excluded in order to explore how variations affect conductivity values. However, both should be considered when analysing results to accurately determine their impact. Therefore, an accurate description of participants should be provided in order to appropriately group them for analysis. In the case no information is provided, it should be made clear participants are assumed to be healthy adults. |
| Yes - 1 | At minimum, participant pathology was provided. Age is further included in the current meta-analysis if available. |
| No - 0 | No information on participant pathology or demographics were provided, or sufficient to assume participants were healthy adults. |
| Unclear – 0.5 | When no detailed description was available, it is still sufficient to assume participants were healthy adults. |
| **Item 2: Does the measurement method appropriately determine the desired value?** |  |
| Pre-specified protocol | The study measures what they set out to measure (i.e. conductivity/resistivity) and reports the values appropriately. |
| Yes - 1 | Results were reported in accordance to the study aims and methodology. |
| No – 0 | Reported values were inconsistent with the study aims and what they claimed to report. |
| Unclear – 0.5 | It is unclear whether or not reported results were in line with the studies aim, therefore an average of 0.5 for this item is assigned. |
| **Item 3: Is the methodology accurately described such that it can be replicated?** |  |
| Pre-specified protocol | Employed methodology should be sufficiently described in order to accurately assess variation in conductivity values due to methodological differences. Enough information should therefore, at minimum, be provided to assign methodology. In order to further validate the chosen method, a sufficient description should allow replication of the method. |
| Yes - 1 | Sufficient information was provided to assign a methodology and replicate their chosen method. |
| No - 0 | No information was provided, and the review is unable to appropriately assign chosen methodology. |
| Unclear 0.5 | It is unclear whether enough information is provided for replication, but methodology can be assigned appropriately. |
| **Item 4: Is the chosen methodology justified?** |  |
| Pre-specified protocol | As there are many different methods measuring conductivity, it would be expected each study would justify their chosen methodology, potentially based on the previous literature. |
| Yes - 1 | Justification for the chosen method was provided. |
| No - 0 | No justification for the employed methodology was provided. |
| Unclear – 0.5 | It is unclear whether or not justification was provided, therefore an average of 0.5 for this item is assigned. |
| **Item 5: Was the measurement protocol verified for accuracy?** |  |
| Pre-specified protocol | Tests measuring conductivity do not come without errors, in order to improve validity of the method, measurement errors should be provided to further verify their results. This may be through simulation or phantom experiments, or as standard deviations within each participant. |
| Yes - 1 | The error associated with the chosen methodology was reported – from simulation, phantom or participant data. |
| No - 0 | No protocol errors were reported. |
| Unclear - 0.5 | It is unclear whether errors were reported or not – i.e. unclear whether reported errors were due to simulation, phantom or participant data errors. |
| **Item 6: Did the measurement and verification method remain unchanged throughout the study?** |  |
| Pre-specified protocol | Methodology should remain constant throughout the measurement process, with any deviations described and justified. |
| Yes - 1 | All measurements were obtained using the same methodology. |
| No – 0 | Methodology did not remain constant. |
| Unclear – 0.5 | It is unclear whether or not the methodology was consistent for each measurement, therefore an average of 0.5 for this item is assigned. |
| **Item 7: Were the measurement errors within an acceptable range? Higher weight?** |  |
| Pre-specified protocol | Errors should be of an acceptable value otherwise use of the chosen method is not justified. Where errors are considerably large, a discussion should be made as to why and ramifications implemented. |
| Yes - 1 | Errors were relatively low and within an acceptable range. |
| No - 0 | Errors were considerably large, and no justifications/ramifications were made. |
| Unclear - 0.5 | No clear error values were provided, or justification/ramification are absent/arbitrary, therefore an average of 0.5 for this item is assigned. |

### Direct Measurement

|  |  |
| --- | --- |
| **Item 8: Were measurements obtained immediately after tissue was excised? If no, how much time elapsed between excision and test?** |  |
| Pre-specified protocol | For conductivity values that reflect the most natural circumstances, measurements should be made immediately after tissue has been excised, or immediately after death. Time-elapsed from excision and measurements may affect the relative conductivity of the tissue. |
| Yes - 1<br>No:<br>< 24 hours – 0.8<br>1 - 7 days – 0.6<br>1 - 8 weeks – 0.4<br>>2 months – 0.2<br>Unclear – 0.5 | No time elapsed from excision and test.<br>Time elapsed between excision and test within the range described and an item score was assigned accordingly.<br>It was unclear how much time elapsed, therefore an average of 0.5 for this item is assigned. |
| **Item 9: Were excised tissue samples kept in 100% saline? If no, what liquid (and concentration) were tissue samples kept in?** |  |
| Pre-specified protocol<br>Yes - 1<br>No:<br>>90% saline – 0.8<br>50-90% saline – 0.6<br><50% saline – 0.4<br>Dry – 0.2<br>Unclear – 0.5 | To ensure conditions are kept as natural as possible, excised tissue should be contained within 100% saline, where conductivity is 1 S/m.<br>Excised tissue samples were contained in 100% saline of 1 S/m conductivity.<br>Tissue samples were kept in varying concentrations of saline and quality scores were assigned accordingly.<br>It is unclear what concentration of saline tissues were kept in, hence they were assigned an average score of 0.5 |
| **Item 10: Was the tissue kept at body temperature (~37 degrees Celsius)?** |  |
| Pre-specified protocol<br>Yes - 1<br>No - 0<br>Unclear – 0.5 | For conductivity values that reflect the most natural circumstances, excised tissue should be maintained at body temperature.<br>Tissue kept at approximately body temperature (34-39°C)<br>Tissue not at body temperature.<br>The temperature of the tissue during conductivity measurements was unclear or ambiguous, therefore an average of 0.5 for this item was assigned. |

### Model-dependent Measurements

|  |  |
| --- | --- |
| **Item 11: Is an individualised head model considered for each participant?** |  |
| Pre-specified protocol<br>Yes, individualistic - 1<br>No, realistic – 0.75<br>No, spherical – 0.25<br>Unclear – 0.5 | Head shape and tissue thickness varies considerably between individuals; therefore, head models should be personalised for each individual – i.e. from MRI/DTI data. If not, realistic head models should be employed as opposed to a spherical model, as the head is not a simplistic sphere.<br>Individualistic head models were considered.<br>A realistic head model was employed.<br>A spherical head model was employed.<br>The head model used was unclear, hence an average quality score of 0.5 is given. |
| **Item 12: Has the head been maximally segmented into appropriate layers depending on the tissue type being measured?** |  |
| Pre-specified protocol<br>**12a: Scalp**<br>Yes: muscle, fat skin- 1<br>Yes: 2 layers - 0.5<br>No - 0<br>N/A<br>**12b: Skull**<br>Yes: 4 layers – 1<br>Yes: 3 layers – 0.66<br>Yes: 2 layers – 0.33<br>No: 1 layer – 0<br>N/A<br>**12c: Brain**<br>Yes: GM + WM – 1<br>No: homogenous – 0.5<br>No: scalp = brain – 0<br>N/A<br>**12d: WM anisotropy**<br>Yes: anisotropic – 1<br>No- isotropic – 0<br>N/A | As specified by sub-sections:<br>The scalp was segmented into the 3 tissue layers of muscle, fat, skin<br>The scalp is considered as 2 layers (i.e. muscle + fat)<br>Scalp is considered one homogenous layer.<br>If the study did not measure scalp conductivity, item 12a is not included in the Quality Assessment score.<br>The skull was segmented into spongiform, inner and outer compact bone with sutures.<br>Skull considered tri-layered; spongiform, inner and outer compact bone without sutures.<br>Skull segmented into spongiform and compact bone.<br>The skull was considered as one homogenous head layer.<br>If the study did not measure scalp conductivity, item 12b is not included in the Quality Assessment score.<br>The brain was compartmented into grey and white matter.<br>The brain was considered as a homogenous tissue.<br>The brain was assumed to have the same conductivity as the scalp.<br>If the study did not measure brain conductivity, item 12c is not included in the Quality Assessment score.<br>WM was modelled as anisotropic<br>WM was modelled as isotropic, or this was not mentioned in the study, therefore WM was assumed to be modelled as isotropic<br>If the study did not measure WM conductivity, item 12d is not included in the Quality Assessment score. |
| **Item 13: Were no assumptions made for the conductivity value of any tissue type?** |  |
| Pre-specified protocol<br>Yes - 1<br>No – 0 | Conductivity values for all reported tissue types were empirically measured rather than assumed from prior literature (i.e. CSF is often assumed to model the remaining tissues).<br>No assumptions were made<br>Conductivity was assumed for one or more tissue types. |

### Model-independent Measurements

|  |  |
| --- | --- |
| **Item 14: Was the magnetic resonance resolution high?** |  |
| Pre-specified protocol<br>Yes: <1mm - 1<br>No: 1-2mm – 0.8<br>No: 2-3mm – 0.6 | High resolution magnetic resonance imaging data with small voxel sizes should be acquired in order to most accurately segment head tissue and improve spatial resolution.<br>MR resolution is 1mm<sup>3</sup> or less.<br>Resolution between 1-2mm<sup>3</sup>.<br>Resolution between 2-3mm<sup>3</sup>. |
| No: 3-4mm – 0.4<br>No: >4mm – 0.2<br>Unclear – 0.5<br>N/A | Resolution between 3-4mm<sup>3</sup>.<br>Resolution above 4mm<sup>3</sup>.<br>The resolution used was unclear or unprovided, hence an average quality score of 0.5 is given.<br>If the study did not employ a magnetic resonance method, item 14 is not included in the Quality Assessment score. |

#### Example 1 & 2: Direct Measurements

Example 1 & 2: Direct Measurements Baumann *et al.* (1997)
| Item | 1 | 2 | 3 | 4 | 5 | 6 | 7 | 8 | 9 | 10 | 11 | 12a | 12b | 12c | 12d | 13 | 14 |
| --- | --- | --- | --- | --- | --- | --- | --- | --- | --- | --- | --- | --- | --- | --- | --- | --- | --- |
| Score | 1 | 1 | 1 | 1 | 0 | 1 | 0.5 | 0.4 | 0.5 | 1 | N/A | N/A | N/A | N/A | N/A | N/A | N/A |
| Total | 7.4 |  |  |  |  |  |  |  |  |  |  |  |  |  |  |  |  |
| QAS | 0.74 (Score total divided by number of relevant items, in this case 10) |  |  |  |  |  |  |  |  |  |  |  |  |  |  |  |  |

Acktari *et al.* (2000)
| Item | 1 | 2 | 3 | 4 | 5 | 6 | 7 | 8 | 9 | 10 | 11 | 12a | 12b | 12c | 12d | 13 | 14 |
| --- | --- | --- | --- | --- | --- | --- | --- | --- | --- | --- | --- | --- | --- | --- | --- | --- | --- |
| Score | 1 | 1 | 1 | 1 | 1 | 1 | 1 | 0.2 | 0.8 | 1 | N/A | N/A | N/A | N/A | N/A | N/A | N/A |
| Total | 9 |  |  |  |  |  |  |  |  |  |  |  |  |  |  |  |  |
| QAS | 0.9 (Score total divided by number of relevant items, in this case 10) |  |  |  |  |  |  |  |  |  |  |  |  |  |  |  |  |

#### Example 3 & 4: Model Dependent Measurements

Example 3 & 4: Model Dependent Measurements Dannhauer *et al.* (2011) – E/MEG
| Item | 1 | 2 | 3 | 4 | 5 | 6 | 7 | 8 | 9 | 10 | 11 | 12a | 12b | 12c | 12d | 13 | 14 |
| --- | --- | --- | --- | --- | --- | --- | --- | --- | --- | --- | --- | --- | --- | --- | --- | --- | --- |
| Score | 1 | 1 | 1 | 1 | 0 | 1 | 0.5 | N/A | N/A | N/A | 1 | 0 | 0.66 | 1 | N/A | 0 | N/A |
| Total | 8.16 |  |  |  |  |  |  |  |  |  |  |  |  |  |  |  |  |
| QAS | 0.68 (Score total divided by number of relevant items, in this case 12) |  |  |  |  |  |  |  |  |  |  |  |  |  |  |  |  |

Fernández-Corazza *et al.* (2017) – EIT
| Item | 1 | 2 | 3 | 4 | 5 | 6 | 7 | 8 | 9 | 10 | 11 | 12a | 12b | 12c | 12d | 13 | 14 |
| --- | --- | --- | --- | --- | --- | --- | --- | --- | --- | --- | --- | --- | --- | --- | --- | --- | --- |
| Score | 1 | 1 | 1 | 1 | 1 | 1 | 1 | N/A | N/A | N/A | 1 | 0 | 0.33 | 1 | N/A | 0 | N/A |
| Total | 9.33 |  |  |  |  |  |  |  |  |  |  |  |  |  |  |  |  |
| QAS | 0.7775 (Score total divided by number of relevant items, in this case 12) |  |  |  |  |  |  |  |  |  |  |  |  |  |  |  |  |

#### Example 5 & 6: Model Independent Measurements

Example 5 & 6: Model Independent Measurements Rullmann *et al.* (2009) - DTI
| Item | 1 | 2 | 3 | 4 | 5 | 6 | 7 | 8 | 9 | 10 | 11 | 12a | 12b | 12c | 12d | 13 | 14 |
| --- | --- | --- | --- | --- | --- | --- | --- | --- | --- | --- | --- | --- | --- | --- | --- | --- | --- |
| Score | 1 | 1 | 1 | 1 | 1 | 1 | 1 | N/A | N/A | N/A | N/A | N/A | N/A | N/A | N/A | N/A | 1 |
| Total | 8 |  |  |  |  |  |  |  |  |  |  |  |  |  |  |  |  |
| QAS | 1 (Score total divided by number of relevant items, in this case 8) |  |  |  |  |  |  |  |  |  |  |  |  |  |  |  |  |

Voigt *et al.* (2011) - MREIT
| Item | 1 | 2 | 3 | 4 | 5 | 6 | 7 | 8 | 9 | 10 | 11 | 12a | 12b | 12c | 12d | 13 | 14 |
| --- | --- | --- | --- | --- | --- | --- | --- | --- | --- | --- | --- | --- | --- | --- | --- | --- | --- |
| Score | 1 | 1 | 1 | 1 | 1 | 1 | 0.5 | N/A | N/A | N/A | N/A | N/A | N/A | N/A | N/A | N/A | 0.8 |
| Total | 7.3 |  |  |  |  |  |  |  |  |  |  |  |  |  |  |  |  |
| QAS | 0.9125 (Score total divided by number of relevant items, in this case 8) |  |  |  |  |  |  |  |  |  |  |  |  |  |  |  |  |

